# Microglia and astrocytes differentially endocytose exosomes facilitating alpha-Synuclein endolysosomal sorting

**DOI:** 10.1101/2022.08.05.502843

**Authors:** M. Pantazopoulou, A. Alexaki, A. Lamprokostopoulou, A. Delis, A. Coens, R. Melki, S.N. Pagakis, L. Stefanis, K. Vekrellis

**Affiliations:** Biomedical Research Foundation Academy of Athens- BRFAA, Clinical- Experimental Surgery & Translational Research, Athens, Greece; Biomedical Research Foundation Academy of Athens- BRFAA, Centre of Basic Research, Athens, Greece; CEA and Laboratory of Neurodegenerative Diseases, Institut Francois Jacob (MIRCen), CNRS, Fontenay-Aux-Roses cedex, France

**Keywords:** exosomes, microglia, astrocytes, endocytosis, endosomal pathway, alpha-synuclein

## Abstract

Exosomes have emerged as key players in cell-to-cell communication in both physiological and pathological processes in the Central Nervous System (CNS). Thus far, the intracellular pathways involved in uptake and trafficking of exosomes within different cell types of the brain (microglia and astrocytes) are poorly understood. In our study, the endocytic processes and subcellular sorting of exosomes were investigated in primary glial cells, particularly linked with the exosome-associated α-synuclein (α-syn) transmission. Mouse microglia and astrocytic primary cultures were incubated with DiI-stained mouse brain-derived exosomes. The internalization and trafficking pathways were analysed in cells treated with pharmacological reagents that block the major endocytic pathways. Brain-derived exosomes were internalized by both glial cell types; however, uptake was more efficient in microglia than in astrocytes. Colocalization of exosomes with early and late endocytic markers (Rab5, Lamp1) indicated that exosomes are sorted to endolysosomes for subsequent processing. Treatment with Cytochalasin D, that blocks actin-dependent phagocytosis and/or macropinocytosis, inhibited exosome entry into glial cells, whereas treatment with inhibitors that strip off cholesterol from the plasma membrane, induced uptake, however differentially altered endosomal sorting. Exosome-associated fibrillar α-Syn was efficiently internalized and detected in Rab5- and Lamp1-positive compartments within microglia. Our study strongly suggests that exosomes enter glial cells through an actin network-dependent endocytic pathway and are sorted to endolysosomes for subsequent processing. Further, brain-derived exosomes are capable of mediating cell-to-glia transmission of pathological α-Syn that is also targeted to the endosomal pathway, suggesting a possible beneficial role in microglia-mediated clearance of toxic protein aggregates, present in numerous neurodegenerative diseases.

## 1. Introduction

Rapid and effective communication between different cell types in the central nervous system (CNS) is required for numerous functions of the brain, i.e., brain development, neural circuit maturation and homeostasis maintenance. Glial cells, including microglia and astrocytes, provide many protective roles for neurons, from immune surveillance, degradation and antigen transfer to synaptic function and plasticity, as well as neuroprotection (Lizarraga-Valderrama and Sheridan, 2021). Neuroglia communicate with neurons and orchestrate CNS homeostasis through numerous pathways. Extracellular vesicles (EV), containing proteins, lipids and nucleic acids, represent an efficient way to transfer cargoes between cells in a functional manner and are categorized, according to their size and origin, to exosomes, microvesicles and apoptotic bodies (EL Andaloussi et al., 2013). Exosomes are small extracellular nanovesicles with a defined structure, originating from endocytosis, with sizes ranging from 30 to 200 nm (Pegtel and Gould, 2019). The main biological composition of exosomes is a bilayer membrane with a spheroid shape, whose structure is defined by the cell origin and is comprised of lipid elements (Skotland et al., 2019), tetraspanins, integrins, etc. (Andreu and Yáñez-Mó, 2014), attributing specificity in target-cell docking and subsequent internalization. This specific composition allows exosomes to cross lipids, biological membranes, as well as the blood brain barrier (BBB). In the CNS, exosomes transfer biomolecules to neighboring cells maintaining homeostasis through the regulation of numerous cell processes, i.e., apoptosis, cell proliferation, inflammation, anti-inflammation, etc. (Gupta and Pulliam, 2014).

Under pathological conditions, exosomes mediate the transfer of nucleic acids (miRNA, mRNA), lipids and toxic proteins, contributing to the development of various diseases, i.e., neural tumors, multiple sclerosis, cerebrovascular diseases, epilepsy, traumatic brain injury, and neurodegenerative diseases (Zhang et al., 2021). Glia-derived exosomes are implicated in the pathogenesis of several neuroinflammatory and neurodegenerative diseases (Gupta and Pulliam, 2014). In neurodegenerative diseases, exosomes orchestrate the mechanisms of secretion, transmission and degradation of miRNAs and misfolded proteins, i.e., A*β*, tau, α-Synuclein (α-Syn), mHtt, etc. In Parkinson’s disease (PD), pathologic α-Syn can be secreted *via* exosomes (Emmanouilidou et al., 2010), spreading this protein to healthy neurons, astrocytes and microglia which in turn may induce inflammatory reactions and cell death (Lee et al., 2010; Harischandra et al., 2019). These events show that intercellular communication through exosomes plays a pivotal role in the pathogenesis of numerous brain diseases, thus targeting such transmission pathways may represent a viable therapeutic strategy.

Exosomes can induce signalling in recipient cells either by direct interaction with extracellular receptors, or by fusing with the plasma membrane, or by internalization *via* a diversity of pathways (Gurung et al., 2021). Mechanisms of classical endocytosis include Clathrin-mediated endocytosis (CME), Caveolin-dependent endocytosis (CDE), Lipid Raft-mediated endocytosis, Macropinocytosis and Phagocytosis. Cumulative studies have described the endocytic pathways exosomes follow, depending on their origin and target cell (Ginini et al., 2022), though, in brain cells, the mechanism(s) mediating exosome uptake still remain elusive. Fitzner and colleagues demonstrated that oligodendroglia-derived exosomes are internalized by microglia through macropinocytosis (Fitzner et al., 2011). In a simplified view, all extracellular material (biomolecules, viruses, etc.) entering a cell can be sorted to early endosomes, mature to late endosomes and can either follow the recycling path back to the plasma membrane or fuse with lysosomes (Hu et al., 2015). The Rab5 GTPase protein is the main regulator of endosome biogenesis and trafficking (Rink et al., 2005), while Lamp1 is considered to define late endosomes, pre-lysosomal vesicles (Humphries et al., 2011) and lysosomes (Marwaha et al., 2017). Although the fate of internalized exosomes is ill-defined, recent studies have shown that exosomes, akin to viruses, can highjack the endolysosomal pathway and in turn release their content within the cytoplasm (Li et al., 2020; Polanco et al., 2021). Altogether, the endocytic pathway(s) exosomes follow, differs among cells depending on the cell origin of exosomes, their interaction with the recipient cells as well as on the fate of exosome cargo upon internalization.

The present study addresses fundamental neurobiology questions in the field of exosome-mediated intercellular communication in the brain. We assessed the internalization and subcellular trafficking of brain-derived exosomes in mouse primary microglia and astrocytes, and hence the mechanistic pathway(s) essential for efficient cell-to-cell communication. Additionally, we sought to designate the differential processes involved in exosome uptake between the different glial cell types, that ultimately reflect their unique cellular function in the brain. Our results show that exosomes are internalized more efficiently in microglia than in astrocytes. Additionally, we provide valuable information on the role of the distinct endocytic pathway(s) and that of the endolysosomal system in the cell processing of exosomes, defining their cellular fate and destination. The actin-dependent endocytic pathways, phagocytosis and/or macropinocytosis, mediate the uptake of exosomes, followed by subcellular sorting to the endolysosomal pathway for further processing. Finally, we evaluated the role of exosomes in clearing protein aggregates involved in the pathogenesis of numerous neurodegenerative diseases. More specificaly, exosomes were preincubated with fibrillar α-Syn assemblies and their internalization, endocytic sorting as well as ultimate fate were evaluated in microglia. We report here that exosome-associated fibrillar α-Syn was targeted to the endolysosomal pathway for subsequent degradation. Our work unravels important intricacies with respect to the competency of the brain-resident macrophage population in pathological protein clearance and homeostasis maintenance.

## 2. Materials and Methods

### 2.1. Mouse brain exosome isolation

Exosomes were isolated from α-synuclein KO mouse brain hemispheres (CD57BL6/JOIaHsd mice, Harlan Laboratories), after removal of olfactory bulbs, cerebellum and meninges, as previously described (Papadopoulos et al., 2018; Karampetsou et al., 2020). Following dissection, brain tissues were digested with papain solution (20 Units/ml, Worthington), activated by L-cysteine (5.5mM) in Hibernate A for 30 minutes, at 37°C. Tissue was homogenized by adding two volumes of cold Hibernate A solution, and the suspension was passed through a 40-µm cell strainer and a 0.2-µm syringe filter. The filtrate was centrifuged at 300xg (10min, 4°C), and then the supernatant was further centrifuged at 2,000xg (10min, 4 °C), 10,000xg (30min, 4°C), and finally at 100,000×g (70min, 4°C). Following aspiration of the supernatant, the exosome pellet was washed in 22–24 ml of cold phosphate-buffered saline (PBS) and centrifuged again at 100,000 × g (70min, 4 °C). Exosome pellet was then diluted in 1.5 ml of sucrose solution (0.95 M), loaded on a sucrose gradient column, and centrifuged at 200,000 × g (16h, 4°C). Sucrose gradient comprises of seven fractions (2–0.25 M, 1.5ml each). Following centrifugation, all fractions were separated according to the gradient (a–g from the top to the bottom), collected individually, and diluted to PBS up to 22 ml of final volume. Following centrifugation at 100,000 × g (70 min, 4 °C), pellets were resuspended in PBS. Exosome enriched fractions were measured according to acetylolcholinesterase (AchE) assays and Micro-Bradford. The content of exosomal sample was evaluated by the ratio of AchE in ng per μg of total protein. Exosomes were labeled with the lipophilic fluorophore Dil (D3911, Invitrogen). Dil was diluted in 50μl of Methanol, with final concentration of 10μM. The mixture was sonicated for 15 minutes, and a consequent mixture of 1 ml of exosome solution (100 ng/ml) in PBS with 6 µl of the 10 µM stock solution, was prepared and gently agitated at 25°C, for 1h. Then, the mixture was centrifuged at 50,000g, at 4°C for 16 hrs. The pellet containing exosomes was resuspended in PBS and stored at -80°C.

### 2.2 Electron Microscopy

Negatively stained electron micrographs were obtained by adding 5μl of exosomes in PBS solution, onto Formvar-coated 400 mesh copper grids, fixing with 0,5% glutaraldehyde (5μl), and staining with a 2% w/w aqueous uranyl acetate solution (Sigma-Aldrich, USA) for 2min. Grids with stained exosomes were examined in a Philips 420 transmission electron microscope at an acceleration voltage of 60 kV and images were acquired with a Megaview G2 CCD camera (Olympus SIS, Munster,Germany).

### 2.3 Human fibrillar α-Synuclein preparation

Human wild-type α-synuclein was expressed in E. coli BL21 DE3 CodonPlus cells and purified as previously described (Ghee et al., 2005). Monomeric, endotoxin free (Pierce LAL Chromogenic Endotoxin Quantification Ki), α-synuclein (200µM) in 50mMTris–HCl, pH 7.5, 150mMKCl was assembled into fibrils by incubation at 37 °C under continuous shaking in an Eppendorf Thermomixer set at 600 r.p.m. for 7 days (Bousset et al., 2013). The assembly reaction was monitored by thioflavin T binding using a Cary Eclipse Fluorescence Spectrophotometer (Varian Medical Systems Inc) set to excitation and emission wavelengths of 440 nm and 480 nm, respectively and the nature of the fibrillar assemblies was assessed by transmission electron microscopy after negative staining with 1% uranyl acetate (Bousset et al., 2013). The resulting fibrils were centrifuged twice at 15,000 g for 10 min and re-suspended twice in PBS. Their concentration was adjusted to 350μM in PBS. They were next fragmented to an average length of 40–50 nm by sonication for 20 min in 2 mL Eppendorf tubes using a Vial Tweeter powered by an ultrasonic processor UIS250 v (250 W, 2.4 kHz; Hielscher Ultrasonic) (Peelaerts et al., 2015). Fragmented fibrils were aliquoted (6 μL) in 0.5 mL Eppendorf tubes, flash frozen in liquid nitrogen and stored at -80 °C until use.

### 2.4 Cell cultures and pharmacological treatment

Glial cells were extracted from P1-P3 mouse brains (CD57BL6/JOIaHsd mice, Harlan Laboratories). Olfactory bulbs, cerebellum and meninges were carefully removed. Brain hemispheres were dissected in Hank’s Balanced Salt Solution (HBSS). The brains were incubated in Dulbecco’s Modifies Eagle’s Media (DMEM) buffer with DNase (10μg/ml final), Trypsin (T4674 Sigma, 2.5 μl/ml final) and Penicillin/Streptomycin (15140122; Invitrogen) for 30-45 minutes, at 37°C. Trypsin was inactivated by adding DMEM with 10% Fetal Bovine Serum (10,270; Gibco). Cells were isolated by centrifugation at 300xg for 7 minutes, the supernatant was removed, and the pellet was diluted in DMEM with 10% FBS and 1% Pen/Strep. Cells were plated (∼7,5-10×10^6^ cells per flask) on poly-D-lysine coated (PDL) (0.01 mg/ml) T75 flasks. Every 3 days, DMEM was renewed. After 7-9 days of dissection, flasks with mixed glial cells were agitated in orbital incubator (150 rpm, 6h) for microglia detachment. The supernatant was harvested, and microglia cells were plated in 24-well dishes (5×10^5^ cells/well) with poly-D-lysine coated glass coverslips. Astrocytic cells were harvested and plated on 24-well plates with poly-D-lysine coated glass coverslips (3×10^5^ cells/well).

24h prior to treatment, cells were washed, and fresh media was added without FBS. Glia cell cultures were treated with or without exosomes (200ng/ml) for 6h, 24h and 48h (cultures were washed 24h post-incubation). Dynasore (ab120192, Abcam, 80μM), Methyl-β-Cyclodextrin (C4555, Sigma, 250μM) and Cytochalasin D (PHZ1063, Invitrogen, 10μM) inhibitors were added concomitantly with exosomes, and cells were incubated over a time course of 6h and 24h. For the treatments concerning α-Syn PFFs (Figure 10, Sup. Fig. 8), microglia cells were single-treated with 200ng/ml PFFs, 400ng/ml exosomes, or double-treated with the combination (PFF+Exo) for 2h and 6h (cultures were washed 2h post-incubation). 20h pre-treatment, the preparations (PFFs, Exo, PFF+Exo) were preincubated at 37°C.

### 2.5 Immunocytochemistry and Confocal Microscopy

Glia cells were fixed with 3.7% formaldehyde and blocking and permeabilization was performed with 0.1% Triton-X100/3% bovine serum albumin (BSA)/ 2% normal goat serum (NGS)/PBS, for 1h at 25°C. Cells were incubated with primary antibodies over night at 4°C and with secondary antibodies and DAPI for 1 hr at 25°C. Primary antibodies used were rabbit monoclonal Rab5 (abcam, ab18211, 1/1000), rabbit monoclonal Lamp1 (abcam, ab24170, 1/1000), mouse monoclonal α-Tubulin (abcam, ab7291, 1/800), rabbit monoclonal Iba1 (Wako, 019-19741, 1/800), and mouse monoclonal D10 (Santa Cruz, sc-515879, 1/1000).

Leica SPS-II inverted confocal microscope with 63x/1.4NA oil immersion objective was used to obtain fluorescence images. Sampling parameters were set following the guidelines of SVI’s PSF calculator (<u>NyquistCalculator</u>). All images were deconvolved with Huygens Essential version 21.10 (Scientific Volume Imaging, The Netherlands, http://svi.nl), using the workflow module based on image metadata with the default parameters.

### 2.6 Imaris Imaging Software Analysis

As described in Sup. Fig. 1, deconvolved confocal stacks were imported into the Imaris software (Bitplane), version 9.1.2, and segmentation of signal of interest from the respective channels (α-Tubulin, exosomes, nuclei, Rab5/Lamp1) was implemented using the “Surface” module. When necessary, “masked” channels, taking the value of zero at pixels lying outside a previously created surface (their mask) and the value of the original channel of interest otherwise, were created. In particular, to measure total exosomes per cell, puncta of exosomes per cell and the mean volume of the individual puncta, a masked exosome channel was created based on the α-Tubulin surface. Then, by creating an exosome surface based on this masked exosome channel, Imaris automatically evaluates these statistics for every object (exosome) comprising this surface. To measure colocalization of exosomes with Rab5 and Lamp1, following the same procedure (α-Tubulin-masked channels to surface), respective surfaces were created. Subsequently the signal from the three α-Tubulin-masked channels (exosomes, Rab5, Lamp1) was masked again based on the corresponding surfaces. The formed (twice) masked channels were used for colocalization analysis by utilizing the “Coloc” module of Imaris. Manders’ coefficients were exported and colocalization was estimated according to the following equation:

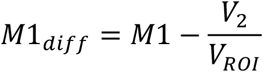

where *M1* is the Manders’ coefficient 1 (percentage of above-threshold signal from channel 1 overlapping with above-threshold signal from channel 2), *V*_*ROI*_ is the total volume of the ROI (region of interest) and *V*_2_ is the volume of the isolated signal from channel 2 (Pike et al., 2017). The colocalization channel and corresponding surface (yellow) were also created. In order to acquire statistics of interest only from internalized exosomes, we used Imaris’ ‘‘Distance transformation’’ function (with the “Distance outside” option) for an exosome surface created from the deconvolved exosome channel (red), computing the distance of the exosomal puncta (original red channel corresponding to exosomes) from the ‘‘α-Tubulin’’ surface. Puncta with value “0” in the resulting “Minimum distance” statistic correspond to exosomes that have entered the cell entirely (called ‘exo inside’) and those in the process of entering (‘exo membrane’). These two groups are characterized by taking zero or positive values, respectively, in the “Maximum distance” statistic also created from the “Distance transformation” function. By selecting and isolating the former puncta into a new surface, their statistics are extracted.

### 2.7 Statistical analysis

GraphPad Prism 9 was used for the statistical analysis. Data were assessed for normality (Shapiro-Wilk test) and two-tailed Student’s *t*-test or Mann-Whitney test was used when comparing two groups, one-way ANOVA with Tukey’s correction for multiple groups, and two-way ANOVA with Tukey’s correction or multiple t-tests for multiple groups with two independent variables. Statistical significance was set as \**p* < .05, \*\**p* < .01, \*\*\**p* < .001, \*\*\*\**p* < .0001 and data were presented as the mean ± SEM from 3 independent experiments, with at least two replicates per assay. No blinding was performed and no test for outliers was conducted.

## 3. Results

### 3.1. Internalization of brain-derived exosomes by Microglia and Astrocytes

To elucidate the internalization process of exosomes in mouse primary microglia and astrocytes, we isolated exosomes from mouse brain by using sequential centrifugations and sucrose gradient ultracentrifugation. Exosomes were quantified by measuring the activity of acetylcholinesterase. Moreover, Electron Microscopy (EM) was used to verify the size, morphology, and structure of exosomes within the isolated fractions (Sup. Figure 1o). In order to follow the internalization of exosomes in primary glial cells, exosomes were prestained with DiI, a fluorescent lipophilic cationic dye. Labelled exosomes (200 ng/ml) were added to primary glia cultures, washed after 24h of incubation, and uptake was monitored 6h, 24h and 48h post-addition. Following immunocytochemistry and confocal microscopy, image analysis was performed using the Imaris Imaging software (Sup. Fig. 1). To evaluate exosome uptake at different time points and conditions, we measured the total volume and number of puncta of exosomes per cell. Values corresponding to the amount of exosomes internalized per cell, as well as the volume of each individual exosomal puncta, revealing the size distribution and intracellular processing, were derived from those measurements. Our data showed that exosomes were internalized by both astrocytes and microglia but at a different rate (Figure 1). As shown in figure 1, after 6h of incubation, microglia cells had uptaken exosomes more efficiently than astrocytes; 7,01 μm^3^ and 4,34 μm^3^ of exosomes per cell, in microglia and astrocytes, respectively. Additionally, total puncta per cell significantly increased after 24h of incubation in microglia but not in astrocytes (Figure 1a,b,c,h,i,j). In microglia cells, the mean volume of the individual puncta remained the same, whereas in astrocytes it increased over time (Figure 1d,k). Next, and in order to distinguish the exosomes that have fully entered the cells from those that were in the process of entering, we used the ‘‘Distance transformation’’ function of the Imaris software, to compute the distance of the exosomal puncta from the ‘‘α-Tubulin’’ surface (Sup. Fig. 1k-n). Hence, we divided exosomes intersecting the cell (‘‘exo total’’) into two separate fractions corresponding to those localized within the cytoplasm (exo inside, henceforth ‘‘exo in’’) and those localized on the plasma membrane (exo membrane, henceforth ‘‘exo mem’’). We measured the proportion of exosomes localized in the cytoplasm (exo in) or on the plasma membrane (exo mem) by calculating the ratio of the exo in or mem (in/mem) volume and number of puncta to total volume and puncta, respectively. Our data suggested that almost 60% of exosomes were internalized (exo in) by microglia, with 40% remaining on the plasma membrane (Figure 1e), whereas in astrocytes this process was slower with more than 70% remaining on the plasma membrane (Figure 1l) after 6h of incubation. 24h post-addition, most exosomes were completely internalized (exo in) by microglia, while they also grew in size, whereas in astrocytes, 40% still resided on the plasma membrane (Figure 1e,l). When the number of exosomal puncta was measured in microglia, 80% of exosomal puncta were internalized (exo in) at 6h (Figure 1f). The puncta localized on the plasma membrane appeared larger than those present inside the cells 6h post-incubation, and their size decreased over time when most of them had fully entered the cell (Figure 1g). In astrocytes, at 24h, 40% of puncta were membrane-associated and much larger in size than those inside. Their size remained the same over time, whereas the size of exosomes localized inside the cells increased (Figure 1m,n), consistent with the overall increase of size of exosomes between 6h and 24h.

**Figure 1.**
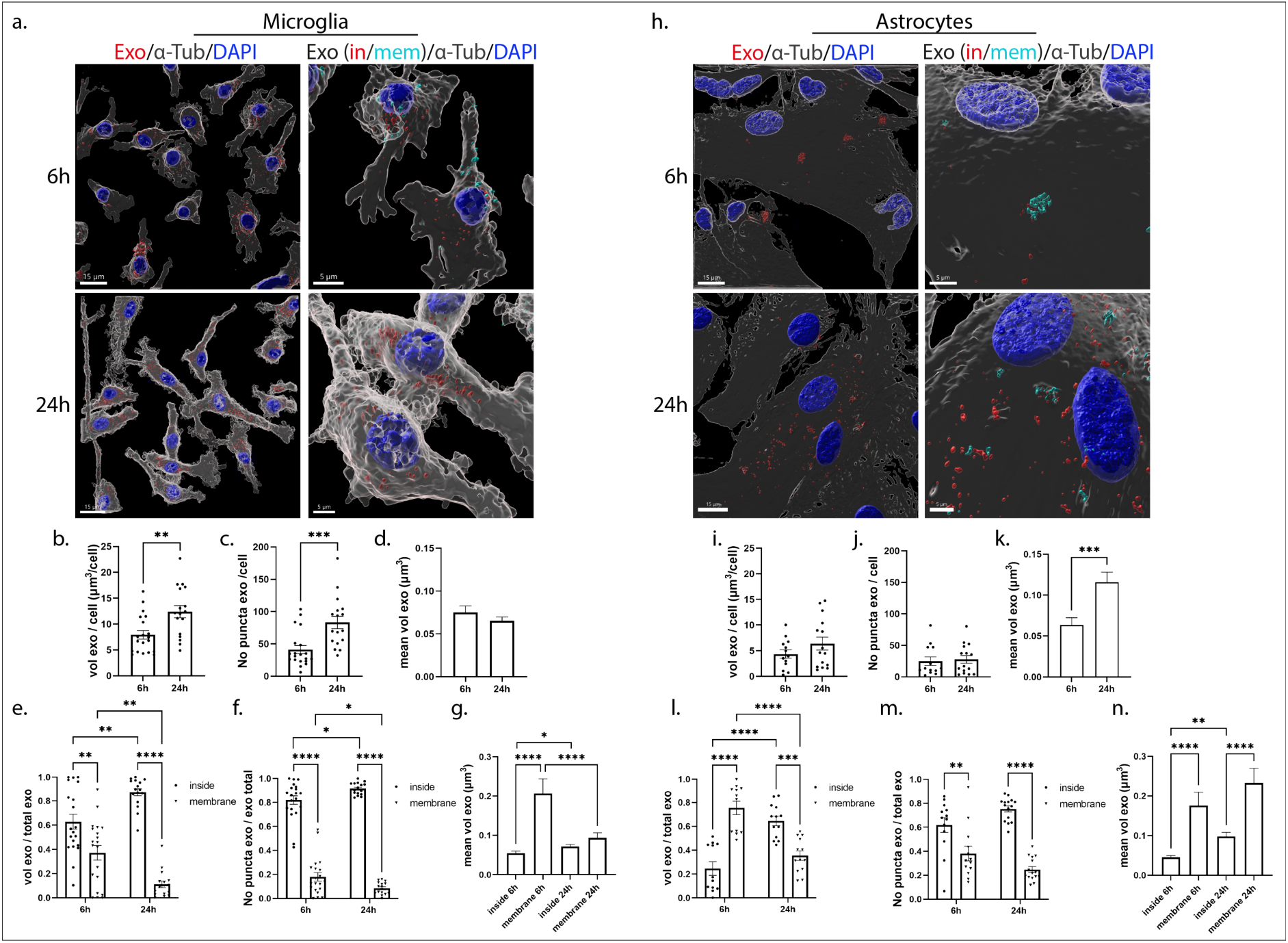
Brain-derived exosomes are internalized in primary microglia and astrocytes. Primary cells were incubated with Dil-stained brain-derived exosomes (depicted in red) for 6h and 24h. Cells were washed, and internalization of exosomes was monitored 6h and 24h post-incubation. Cells were fixed and immunostained with an antibody against α-Tubulin (α-Tub) (gray) while cell nuclei were stained with DAPI (blue). Confocal images were deconvolved and analyzed with the Imaris Imaging software. Exosomes were defined as cytoplasmic (exo in, red) or membranous (exo mem, cyan blue) by using the ‘‘Distance transformation’’ module of the Imaris Imaging software, computing the distance of the exosomal puncta from the ‘‘α-Tubulin’’ surface. Representative Imaris images depict the internalization of exosomes masked with the α-Tubulin surface (left panel, scale bar 15 μm) and in/mem exosomes (right panel, scale bar 5 μm) in microglia (a) and astrocytes (h), 6h and 24h post-addition. Graphs show the total volume of internalized exosomes per cell (b, i), the number of puncta per cell (c, j), the mean volume of exosomes (d, k), the ratio of the volume of exosomes (in/mem) per total volume (e, l), the ratio of the number of puncta (in/mem) per total number (f, m), and the mean volume of exosomes (in/mem) (g, n), in microglia and astrocytes, respectively. Data are presented as the mean ± SEM of minimum 3 independent cell preparations, with at least two replicates per assay; Student’s t-test was used for (b), (d), (i) and (k), Mann-Whitney test for (c) and (j), one-way ANOVA with Tukey’s correction for (g) and (n), two-way ANOVA with Tukey’s correction for (e) and (m) and multiple t-test for (f) and (m). Statistical significance was set as *p < .05, **p < .01, ***p < .001, ****p < .0001.

When the medium was changed after 24h of incubation, the total volume of exosomes remained the same between 24h and 48h post-addition with most exosomes residing inside both glial cell types (Sup. Fig. 2a,b,e,f,h,i,l,m). Since the DiI fluorophore is stable even in acidic environments, no alteration in the total volume was expected. In microglia, the mean volume of the individual puncta increased 48h post-addition, due to the increase in size of the few membranous puncta (Sup. Fig. 2d,g). In astrocytes, the mean volume of exosomes localized on the plasma membrane decreased at 48h (Sup. Fig. 2n) as in microglia after 24h of exosome incubation. Overall, both glial cell types uptake brain-derived exosomes, however, the process is faster in microglia than in astrocytes.

### 3.2. Endocytic trafficking of exosomes in Microglia and Astrocytes

Next, we wished to elucidate the endocytic pathway exosomes follow post-internalization in primary glial cells. Thus, we used two endocytic markers of the early and late endocytic pathway. Rab5 is a key regulator for the formation of early endosomes (EE) (Rink et al., 2005), and LAMP1 (lysosomal-associated membrane protein 1) is associated with the late endosomal, pre-lysosomal and lysosomal pathway (LE/Lysosome) (Humphries et al., 2011; Marwaha et al., 2017). Therefore, we investigated whether exosomes are sorted in EE and LE/Lysosome 6h, 24h and 48h following their addition to cells. To follow the trafficking of exosomes, Dil-labeled exosomes were added to primary glial cultures, washed off after 24h of incubation and the cells were fixed and immunostained with antibodies against Rab5 or Lamp1. We examined the colocalization of exosomes with the endosomal markers, by performing immunofluorescence and confocal microscopy followed by Imaris analysis that permitted 3D reconstruction analysis of pixel colocalization between Dil-stained exosomes and endosomes (EE, LE/Lysosome). Quantitative analysis indicated that in microglia cells approximately 60% of internalized exosomes colocalize with Rab5 at 6h and 40% at 24h post-addition, with the mean volume of colocalized puncta decreasing over time (Figure 2a-c). 24h following washing the cells (48h post-treatment), the percentage of colocalization between exosomes and Rab5 remained the same as did the size of colocalized puncta (Sup. Figure 3a-c). Additionally, 60% of exosomes colocalized with Lamp1, and this percentage did not change over time. The size of the colocalized puncta also remained the same throughout the experimental timeframe (Figure 2d-f, Sup. Fig. 3d-f).

**Figure 2.**
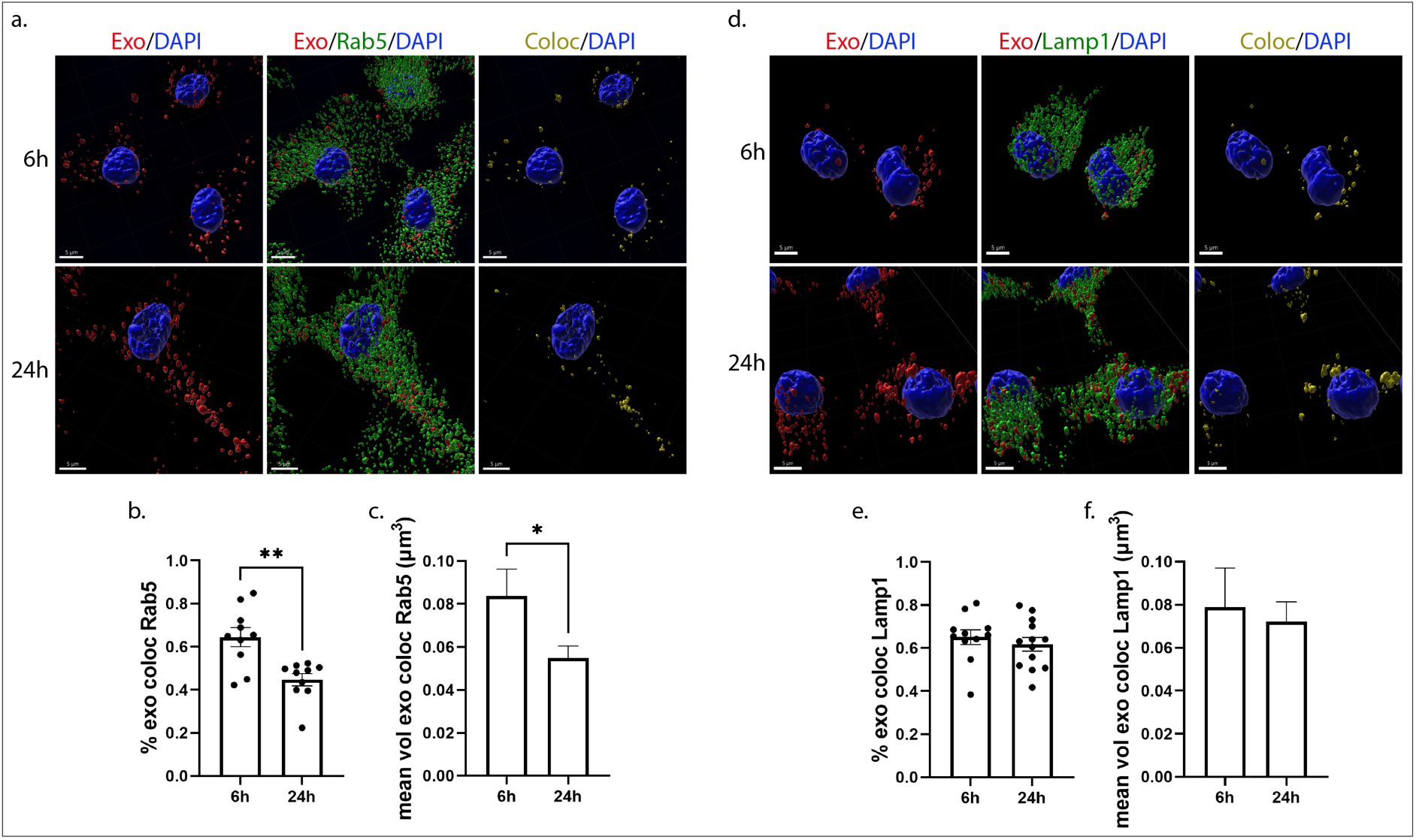
Exosomes when uptaken by primary microglia follow the endocytic pathway and are colocalized with Rab5 (early endosomes-EE) and Lamp1 (late endosomes-LE/Lysosomes). Microglia cells were incubated with Dil-labeled exosomes (red), for 6h and 24h. Colocalization of exosomes with Rab5 and Lamp1 was monitored at 6h and 24h post-treatment. Cells were fixed and immunolabeled for Rab5 or Lamp1 (green), α-Tubulin (α-Tub) (gray) and DAPI (blue), detecting cell nuclei. Representative Imaris images depict colocalization between internalized exosomes and the endocytic markers, Rab5 (a-c) and Lamp1 (d-f), after 6h and 24h of incubation. The colocalization channel and surface (yellow) were built using the Imaris imaging software. Scale bar 5 μm. Graphs show colocalization between exosomes and Rab5/Lamp1 (Manders’ Colocalization Coefficient) after 6h and 24h (b and e) of treatment as well as the mean volume of puncta colocalized with Rab5 (c) and Lamp1 (f) at the different time points. Data are presented as the mean ± SEM of minimum 3 independent cell preparations, with more than 80 cells measured; Student’s t-test was used, and statistical significance was set as *p < .05, **p < .01, ***p < .001, ****p < .0001.

In astrocytes, approximately 80% of exosomes colocalized with Rab5 (Figure 3a,b) and 60-70% with Lamp1 (Figure 3d,e), at 6h and 24h of treatment. The size of colocalized puncta increased over time in both cases as exosomes fused with the early endosomal or late endosomal compartments (Figure 3c,f). In contrast, after washing the cells (48h post-treatment), the percentage of colocalization between exosomes and Rab5 decreased to 60%, as did the volume of colocalized puncta (Sup. Fig. 4a-c). No differences were observed concerning the colocalization of exosomes with LE/lysosomes, however the size of colocalized puncta decreased over time, probably due to fusion processes with the lysosome (Sup. Fig. 4d-f). Our data indicate that exosomes are targeted to the endolysosomal pathway in both glial cell types, however the distribution to EE or to LE/Lysosomes differs when comparing microglia and astrocytes.

**Figure 3.**
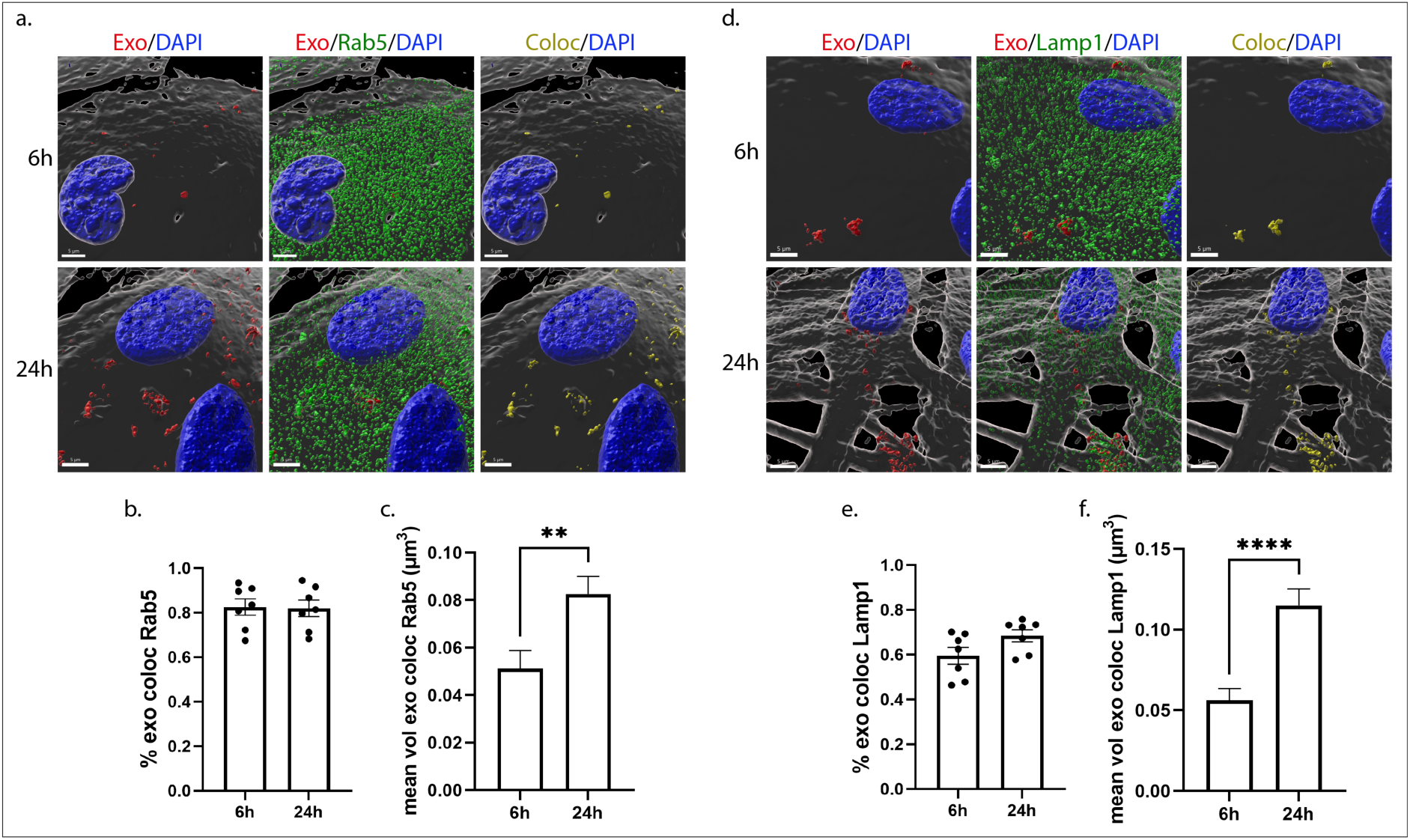
Exosomes are targeted to the endocytic pathway and are colocalized with Rab5 (EE) and Lamp1 (LE/Lysosomes) following addition on primary astrocytes. Cells were incubated with Dil-stained exosomes (red) for 6h and 24h, and colocalization of exosomes with Rab5 and Lamp1 was measured 6h and 24h post-treatment. Cells were fixed and immunostained for Rab5 and Lamp1 (green), α-Tubulin (α-Tub) (gray) and DAPI (blue). Representative Imaris images depict colocalization between internalized exosomes and the endocytic markers, Rab5 (a-c) and Lamp1 (d-f), after 6h and 24h of treatment. The colocalization surface is depicted in yellow. Scale bar 5 μm. Graphs show the colocalization between exosomes and Rab5/Lamp1 (Manders’ Colocalization Coefficient) after 6h and 24h (b and e) of treatment as well as the mean volume of puncta colocalized with Rab5 (c) and Lamp1 (f). Data are presented as the mean ± SEM of minimum 3 independent cell preparations, with more than 60 cells measured; Student’s t-test was used, and statistical significance was set as *p < .05, **p < .01, ***p < .001, ****p < .0001.

### 3.3. Effect of the Dynamin-dependent endocytic pathway inhibition on the uptake and endocytic trafficking of exosomes in primary glia cells

Dynamin is the key regulator of clathrin-and caveolin-dependent endocytosis (Vallee et al., 1993). To assess the role of these pathways in the internalization and intracellular trafficking of exosomes, we used Dynasore (Dyn), a pharmacological reagent that inhibits dynamin (Newton et al., 2006). DiI-labelled exosomes (200 ng/ml) were added to primary cultures in the absence (NT) or presence of Dyn and uptake was monitored at 6h and 24h post-addition. Immunocytochemistry and confocal microscopy, followed by Imaris analysis, showed that exosomes were taken up by microglia and no difference was observed in the total volume of exosomes per cell between NT and Dyn-treated cells (Figure 4a,b). Interestingly, in Dyn-treated cells, the total volume and the number of puncta remained comparable between time points (Figure 4b,c). There was a statistical, albeit slight increase in the mean volume of puncta upon Dyn-treatment, 6h post-incubation (Figure 4d) but no differences were reflected in the in/mem puncta size (Sup. Fig. 5c,d). Likewise, no differences were observed in the proportion of in/mem exosomes (volume and puncta) or in their size between NT and Dyn-treated cells, at any time point (Figure 4e,f, Sup. Fig. 5a-d). After 24h of incubation, the majority of exosomes (volume and puncta) had been efficiently targeted from the plasma membrane to the cytoplasm in both conditions.

**Figure 4.**
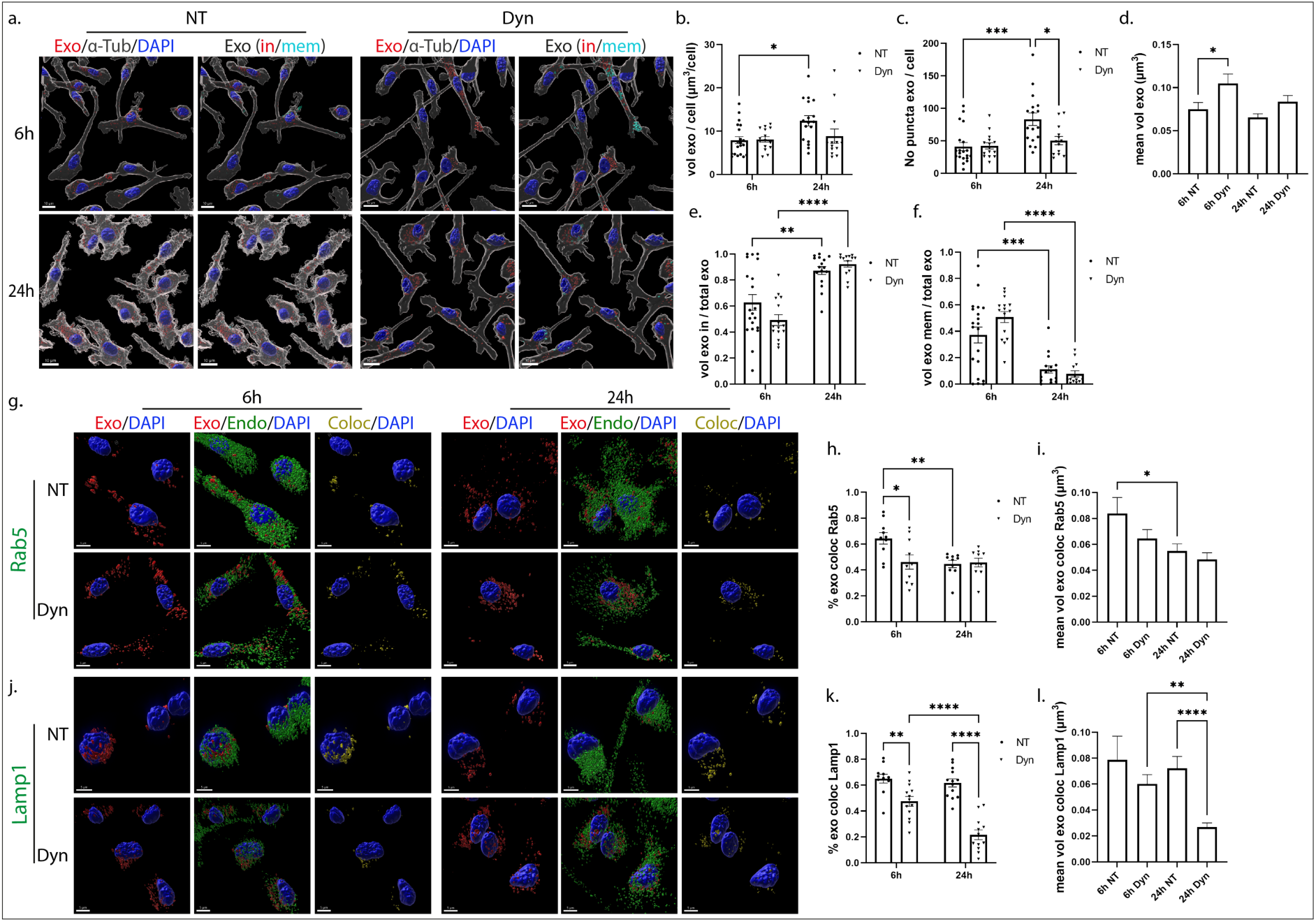
Uptake and endocytic trafficking of exosomes in primary microglia following inhibition of the Dynamin-dependent endocytic pathway. Cells were incubated with Dil-labeled exosomes (red), in the absence (NT) or presence of Dynasore (Dyn) for 6h and 24h. Cells were fixed and immunostained against α-Tubulin (α-Tub) (gray) and DAPI (blue). Representative Imaris images depict the internalization of exosomes masked with the α-Tubulin surface and in/mem exosomes without (NT) or with Dynasore (Dyn) at 6h and 24h (a) of incubation with exosomes. Scale bar 10 μm. Graphs show the total volume of internalized exosomes per cell (b), the number of exosomal puncta per cell (c), the mean volume of internalized exosomes (d), the percentage of exosomal volume, inside (e) or on the membrane (f) per total volume of exosomes. The endocytic trafficking of exosomes was evaluated by measuring the colocalization of exosomes with Rab5 (g-i) and Lamp1 (j-l) Scale bar 5 μm. Graphs show colocalization between exosomes and Rab5 (h) or Lamp1 (k) and the mean volume of colocalized puncta (i and l, respectively) at different time points. Data are presented as the mean ± SEM of minimum 3 independent cell preparations, with at least two replicates per assay; one-way ANOVA with Tukey’s correction was used for (d), (i) and (l), two-way ANOVA with Tukey’s correction for (b), (e), (f), (h) and (k) and multiple t-test for (c). Statistical significance was set as *p < .05, **p < .01, ***p < .001, ****p < .0001.

To gain further insight into the endocytic trafficking of exosomes when dynamin is inhibited, colocalization with EE and LE/Lysosomes was assessed. Our data indicate that colocalization of exosomes with Rab5 (Figure 4g-i) and Lamp1 (Figure 4j-l) decreased approximately 20% following Dyn treatment at 6h. The percentage of colocalization with Lamp1 and the size of Lamp1-colocalized puncta decreased over time. Altogether, exosomes enter microglia in a clathrin- and caveolin-independent manner, while their targeting to and processing through the endolysosomal pathway is partially hindered upon Dyn treatment.

In astrocytes, Imaris analysis indicated that upon Dyn treatment, the total volume of exosomes internalized was the same as in control NT conditions. However, increased number of internalized puncta (exo in) were detected after 6h of incubation (Figure 5a-c). There was a statistically significant increase in the mean volume of puncta over time in NT and Dyn treated cells within 24h (Figure 5d). Comparing the fractions, in/mem, between the treatments, we observed that less exosomes were confined on the plasma membrane in Dyn-treated cells (ratio of volume and puncta), with more than 60% residing within the cytoplasm, 6h post-addition (Figure 5e,f, Sup. Fig. 5e,f). The size of membrane-associated puncta is smaller at both time points and those inside remain the same over time compared to control NT cells (Sup. Fig. 5g,h). Assessing the intracellular distribution of exosomes in the endocytic compartments, it appears that exosomes escape the endocytic pathway to a great extent as evidenced from the 60% and 40% decrease in exosome colocalization with Rab5 and Lamp1, respectively (Figure 5g-l). Additionally, the size of Rab5-colocalized puncta increased overtime, in a treatment-independent manner, and that of Lamp1-colocalized puncta remained unaltered overtime in Dyn-treated cells. Our data suggest that in Dyn-treated astrocytes internalization of exosomes was faster at early time points, with less exosomes residing on the plasma membrane, whereas no effect was depicted in microglia. In both cell types, exosomes failed to a greater or lesser extent to fuse with EE and LE/Lysosomes.

**Figure 5.**
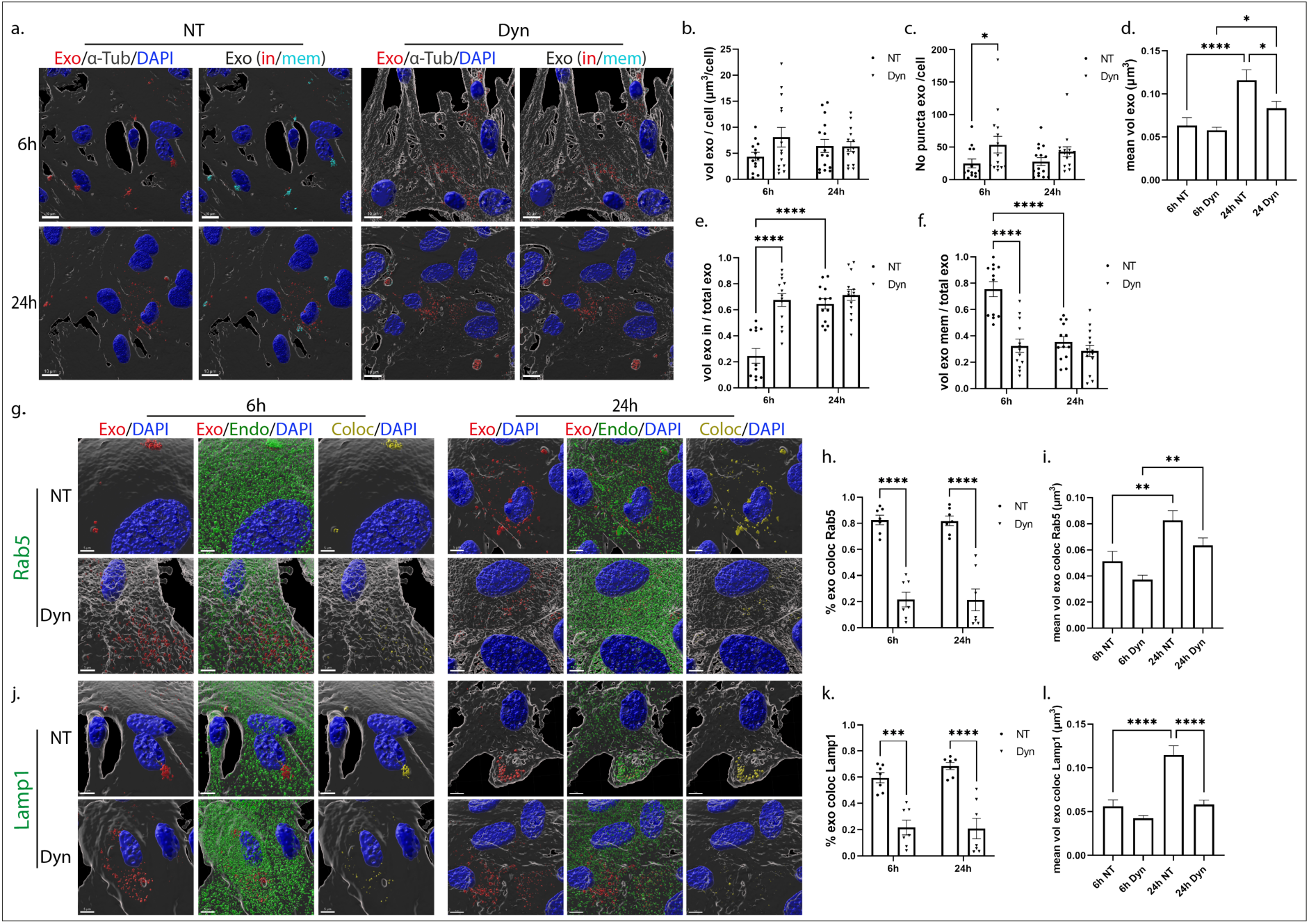
Internalization and endocytic trafficking of exosomes in primary astrocytes following Dynasore treatment. Cells, incubated with Dil-labeled exosomes (red), in the absence (NT) or presence of Dynasore (Dyn) for 6h and 24h, were fixed and immunostained against α-Tubulin (α-Tub) (gray) and DAPI (blue). Representative Imaris images depict the internalization of exosomes masked with the α-Tub surface and in/mem exosomes in cells treated without or with Dyn at 6h and 24h (a). Scale bar 10 μm. Graphs show the total volume of internalized exosomes per cell (b), the number of exosomal puncta per cell (c), the mean volume of internalized exosomes (d), the percentage of exosomal volume, inside (e) or membrane (f) per total volume, after 6h and 24h of treatment. The endocytic trafficking of exosomes was monitored by measuring the colocalization of exosomes with Rab5 (g-i) and Lamp1 (j-l). Scale bar 5 μm. Graphs show the colocalization between exosomes and Rab5 (h) or Lamp1 (k) and the mean volume of colocalized puncta (i and l, respectively) at different time points. Data are presented as the mean ± SEM of minimum 3 independent cell preparations, with at least two replicates per assay; one-way ANOVA with Tukey’s correction was used for (d), (i) and (l), two-way ANOVA with Tukey’s correction for (b), (e), (f), (h) and (k) and multiple t-test for (c). Statistical significance was set as *p < .05, **p < .01, ***p < .001, ****p < .0001.

### 3.4. Uptake and endocytic trafficking of exosomes in primary glia cells following inhibition of Lipid Raft-mediated endocytosis

Lipid rafts are defined as micro-domains within the plasma membrane, enriched in cholesterol and sphingolipids, participating in endocytic processes further classified as caveolae- and dynamin-dependent endocytosis, or non-caveolae pathways (dynamin-dependent or independent) (El-Sayed and Harashima, 2013). Methyl-β-cyclodextrin (Cyclo) is an extensively used pharmacological inhibitor of the pathway, inducing depletion of cholesterol from the plasma membrane (Dutta and Donaldson, 2012). In our experiments, glial cells were incubated with DiI-stained exosomes, with or without methyl-β-cyclodextrin, for 6h and 24h. Imaris analysis demonstrated that in microglia, upon methyl-β-cyclodextrin inhibition, exosomes entered cells faster at early time points (Figure 6a,b), whereas no difference was detected in the total number of puncta (Figure 6c) or in the proportions of in/mem exosomes (volume and number of puncta) (Figure 6e,f, Sup. Fig. 6a,b). Regarding the size of internalized exosomes, exosomal puncta were larger at early time points, mirroring the size of exosomes of the inside fraction, which seemed to be larger in size at both time points compared to control NT conditions (Figure 6d, Sup. Fig 6c,d), probably due to the increased internalization rate. Staining with endosomal markers demonstrated that exosomes reside in EE to a great extent, reaching 60% colocalization with Rab5 even after 24h of incubation (Figure 6g-i) whereas no differences were detected concerning colocalization with Lamp1 (Figure 6j-l). The size of endosome-colocalized puncta is the same with that of the NT cells, at both time points.

**Figure 6.**
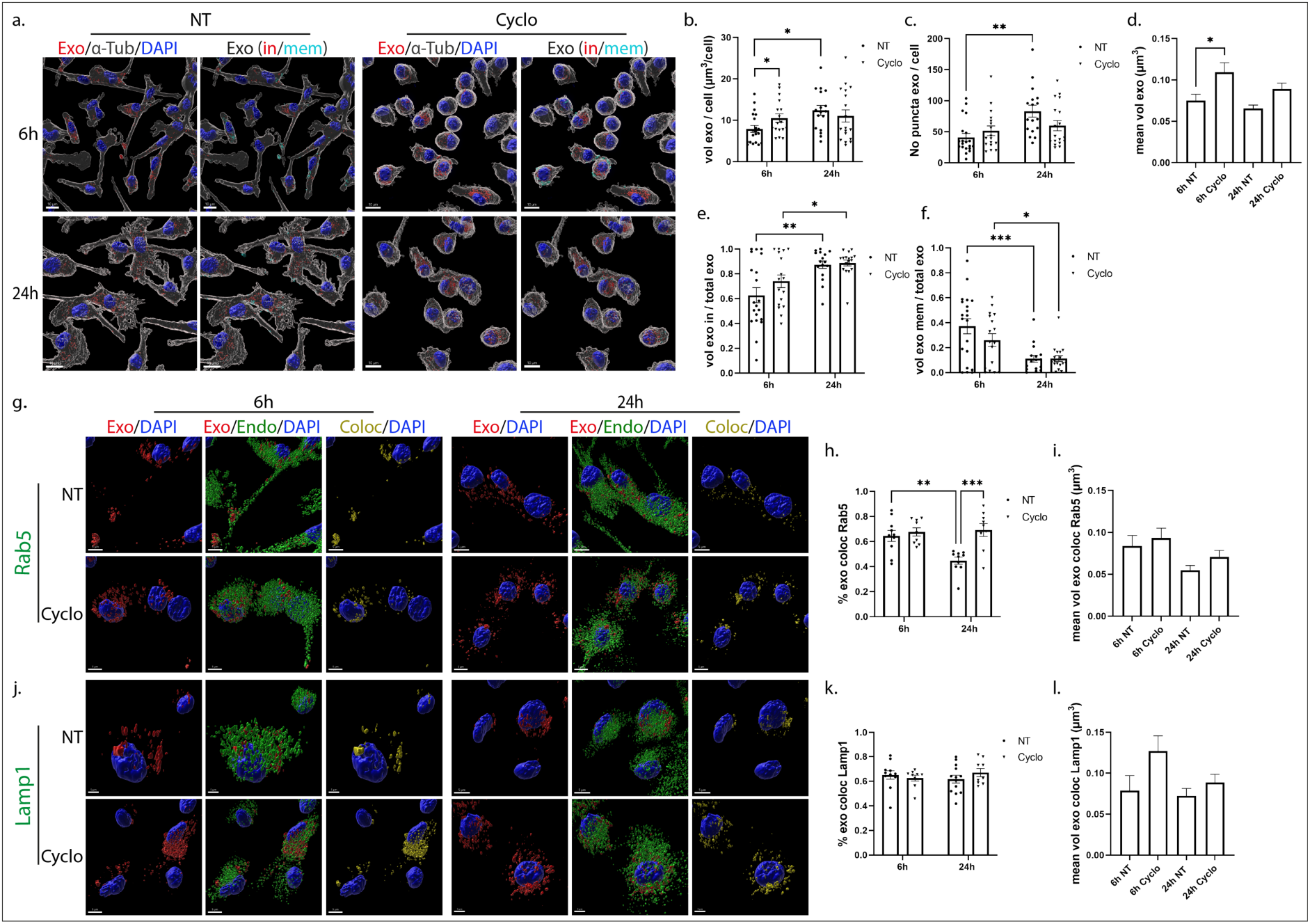
Uptake and endocytic trafficking of exosomes in primary microglia upon depletion of cholesterol with methyl-β-cyclodextrin. Cells, treated with Dil-labeled exosomes (red), in the absence (NT) or presence of methyl-β-cyclodextrin (Cyclo) for 6h and 24h, were fixed and immunostained against α-Tubulin (α-Tub) (gray) and DAPI (blue). Representative Imaris images depict the internalization of exosomes and in/mem exosomes without or with Cyclo at 6h and 24h (a) of incubation. Scale bar 10 μm. Graphs show the total volume of internalized exosomes per cell (b), the number of exosomal puncta per cell (c), the mean volume of internalized exosomes (d), the percentage of exosomal volume, inside (e) or membrane (f) per total volume, 6h and 24h post-incubation. Colocalization of exosomes with Rab5 (g-i) and Lamp1 (j-l) was measured. Scale bar 5 μm. Graphs show colocalization between exosomes and Rab5 (h) or Lamp1 (k) and the mean volume of colocalized puncta (i and l, respectively) at different time points, with or without Cyclo. Data are presented as the mean ± SEM of minimum 3 independent cell preparations, with at least two replicates per assay; one-way ANOVA with Tukey’s correction was used for (d), (i) and (l), two-way ANOVA with Tukey’s correction for (e), (f), (h) and (k) and multiple t-test for (b) and (c). Statistical significance was set as *p < .05, **p < .01, ***p < .001, ****p < .0001.

In primary astrocytic cells, methyl-β-cyclodextrin treatment demonstrated a more prominent effect, with increased total volume and number of puncta 6h post-addition. Additionally, the proportion of inside exosomes also increased, reaching more than 70%, compared to control NT conditions, where only 20% had fully entered the cells (Figure 7a,b,c,e,f, Sup. Fig. 6e,f). Surprisingly, 24h post-incubation the total volume and the number of puncta per cell decreased, reaching values similar to the ones observed in NT cells, while the size of exosomes (total, in and mem) remained unaltered over time, in cyclo-treated cells (Figure 7d, Sup. Fig. 6g,h). A number of studies link cholesterol with endosomal flux and endosome fusion and fission dynamics (Boura et al., 2012; Glitscher and Hildt, 2021). Monitoring the endocytic trafficking in cyclo-treated cells, we noticed that internalized exosomes failed to reside within EE, with almost 40% decrease (Figure 7g-i) at both time points, while Lamp1-positive vesicles were mostly present at later time points (Figure 7j-l). Altogether, our data suggest that exosomes enter astrocytes faster when cholesterol is extracted from the plasma membrane and are targeted directly to the lysosome, thus exhibiting a decreased size (Figure 7d) after fusion with lysosomes over time.

**Figure 7.**
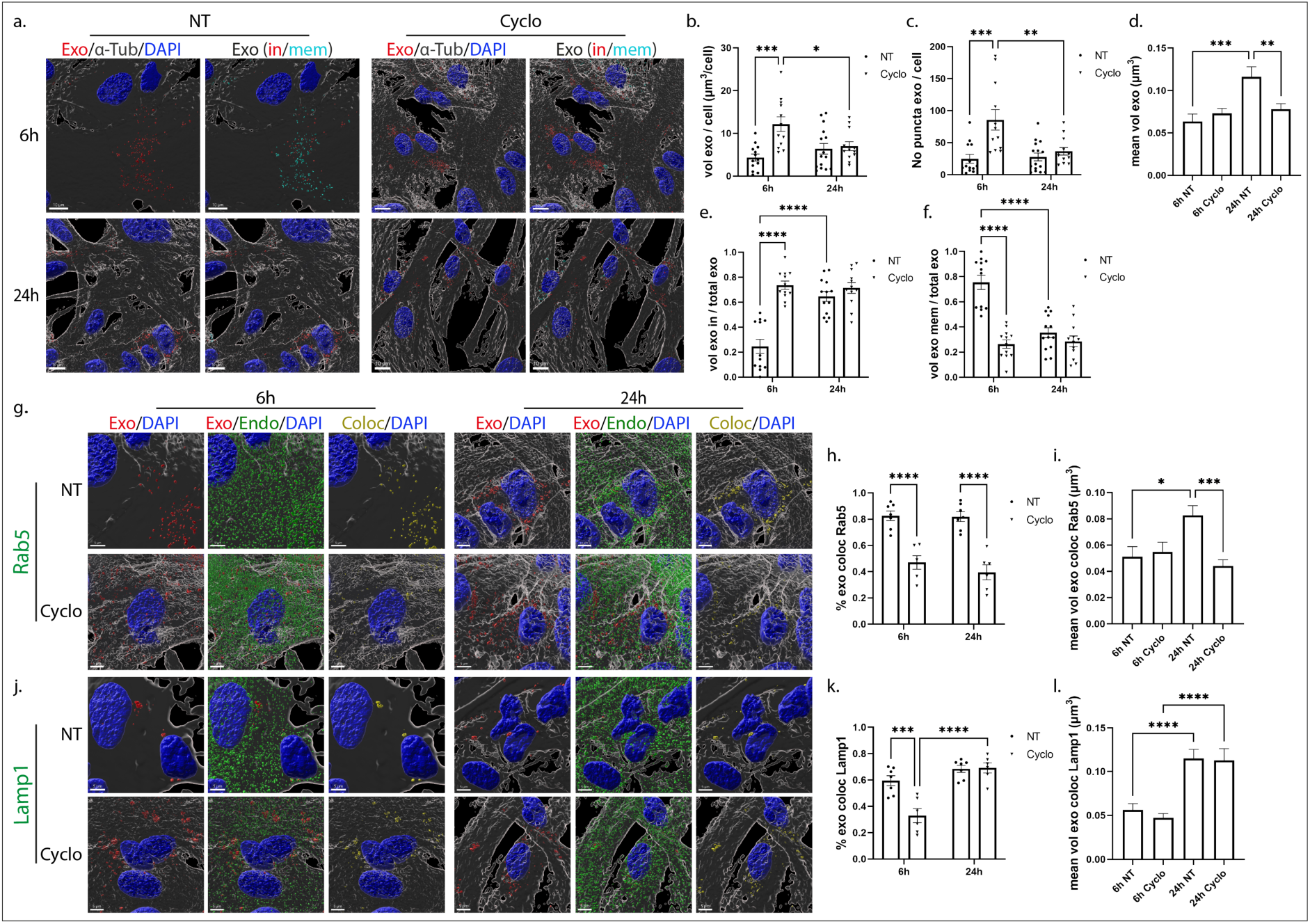
Internalization and endocytic trafficking of exosomes in primary astrocytes upon methyl-β-cyclodextrin (Cyclo) treatment. Cells, treated with Dil-labeled exosomes (red), in the absence (NT) or presence of methyl-β-cyclodextrin (Cyclo) for 6h and 24h, were fixed and immunostained against α-Tubulin (α-Tub) (gray) and DAPI (blue). Representative Imaris images show the internalization of exosomes and in/mem exosomes without or with Cyclo 6h and 24h (a) post-treatment. Scale bar 10 μm. Graphs present the total volume of internalized exosomes per cell (b), the number of exosomal puncta per cell (c), the mean volume of internalized exosomes (d), the percentage of exosomal volume, inside (e) or membrane (f) per total volume, 6h and 24h post-incubation. Colocalization of exosomes with Rab5 (g-i) and Lamp1 (j-l) was monitored. Scale bar 5 μm. Graphs show colocalization between exosomes and Rab5 (h) or Lamp1 (k) and the mean volume of colocalized puncta (i and l) at different time points, with or without Cyclo. Data are presented as the mean ± SEM of minimum 3 independent cell preparations, with at least two replicates per assay; one-way ANOVA with Tukey’s correction was used for (d), (i) and (l), two-way ANOVA with Tukey’s correction for (b), (e), (f), (h) and (k) and multiple t-test for (c). Statistical significance was set as *p < .05, **p < .01, ***p < .001, ****p < .0001.

### 3.5. The role of actin-dependent endocytic pathway on the internalization and endocytic trafficking of exosomes in primary glia cells

To inhibit the pathways of phagocytosis and macropinocytosis, we used the pharmacological reagent Cytochalasin D, that disrupts actin network organization, indispensable for both endocytic pathways (Ginini, 2022). DiI-labeled brain derived exosomes were added on glial cells, in the absence or presence of Cytochalasin D (Cyto), for 6h and 24h. Analysis with the Imaris Imaging Software demonstrated compromised exosome entry into microglia cell cytoplasm upon actin-dependent endocytic pathways inhibition as assessed by measurements of total volume and number of puncta per cell as well as from the proportions of the in/mem fractions (volume, number of puncta) (Figure 8a,b,c,e,f, Sup. Fig. 7a,b). Upon Cyto-treatment, more than 90% of exosomes accumulated on the plasma membrane at both time points, creating large puncta unable to enter cells (Figure 8d,e,f, Sup. Fig. 7c,d). Analysis of the endocytic trafficking showed increased colocalization with Rab5, 70-80%, resulting in larger colocalized puncta at both time points (Figure 8g-i). Additionally, similar to what we observed in control NT cells, exosomes that entered cells resided in Lamp1-positive compartments after 6h and 24h of incubation, however their size increased in Cyto-treated cells, 6h post-incubation (Figure 8j-l).

**Figure 8.**
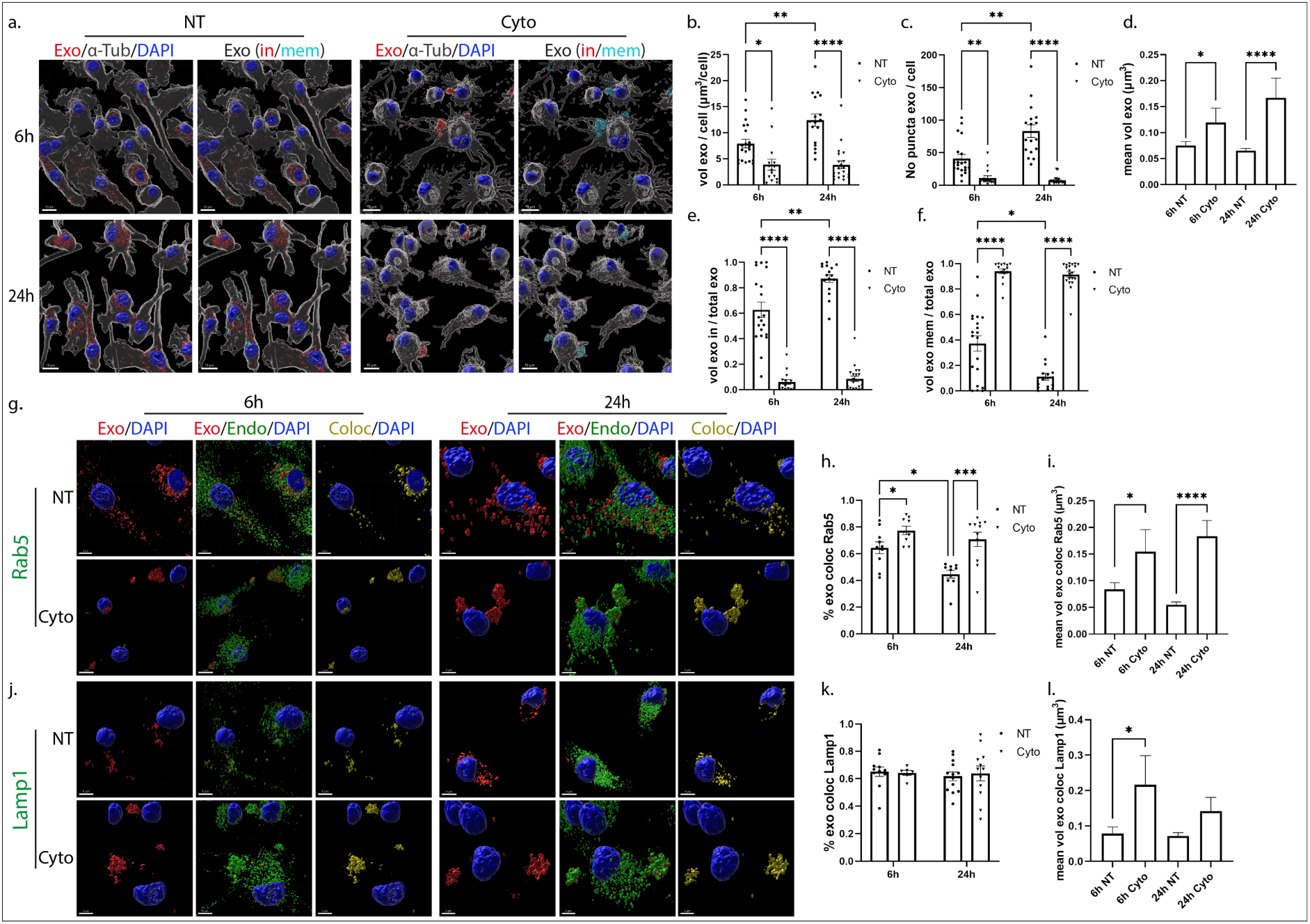
Internalization of exosomes in primary microglia following inhibition of the actin-dependent endocytic pathway. Cells were incubated with Dil-labeled exosomes (red), in the absence or presence of Cytochalasin D (Cyto) for 6h and 24h. Cells were fixed and immunostained against α-Tubulin (α-Tub) (gray) and nuclei were stained with DAPI (blue). Representative Imaris images show the internalization of exosomes and in/mem exosomes without or with Cyto, 6h and 24h (a) post-treatment. Scale bar 10 μm. Graphs present the total volume of internalized exosomes per cell (b), the number of exosomal puncta per cell (c), the mean volume of internalized exosomes (d), the percentage of exosomal volume, inside (e) or membrane (f), per total volume, 6h and 24h post-incubation. Colocalization of exosomes with Rab5 (g-i) and Lamp1 (j-l) was monitored. Scale bar 5 μm. Graphs show colocalization between exosomes and Rab5 (h) or Lamp1 (k) and the mean volume of colocalized puncta (i and l, respectively) at different time points, with or without Cyto. Data are presented as the mean ± SEM of minimum 3 independent cell preparations, with at least two replicates per assay; one-way ANOVA with Tukey’s correction was used for (d), (i) and (l), two-way ANOVA with Tukey’s correction for (h) and (k) and multiple t-test for (b), (c), (e) and (f). Statistical significance was set as *p < .05, **p < .01, ***p < .001, ****p < .0001.

Likewise, in astrocytes, inhibition of the actin-dependent pathways inhibited exosome entry with over 90% remaining on the plasma membrane at both time points (Figure 9a,b,c,e,f, Sup. Fig. 7e,f). Even though the size of total puncta did not change upon Cyto treatment, the differences in puncta size of the mem fraction between the treatments were more prominent 24h post-incubation, a difference not observed at 6h of incubation, as exosomes still resided on the plasma membrane in NT cells (Figure 9d, Sup. Fig. 7g,h). Even though exosomes were restricted to the plasma membrane, the components of the endocytic trafficking (Rab5 and Lamp1) were found colocalized with them on the plasma membrane (Figure 9g,h,j,k), possibly attempting to process those that are blocked from entering cells. Changes in the size of Rab5-colocalized puncta were detected 6h post-incubation and of Lamp1-colocalized puncta after 24h (Figure 9i,l). Altogether, the actin-dependent endocytic pathways, phagocytosis and/or macropinocytosis, seemed to be responsible for the internalization of exosomes in both glial cell types.

**Figure 9.**
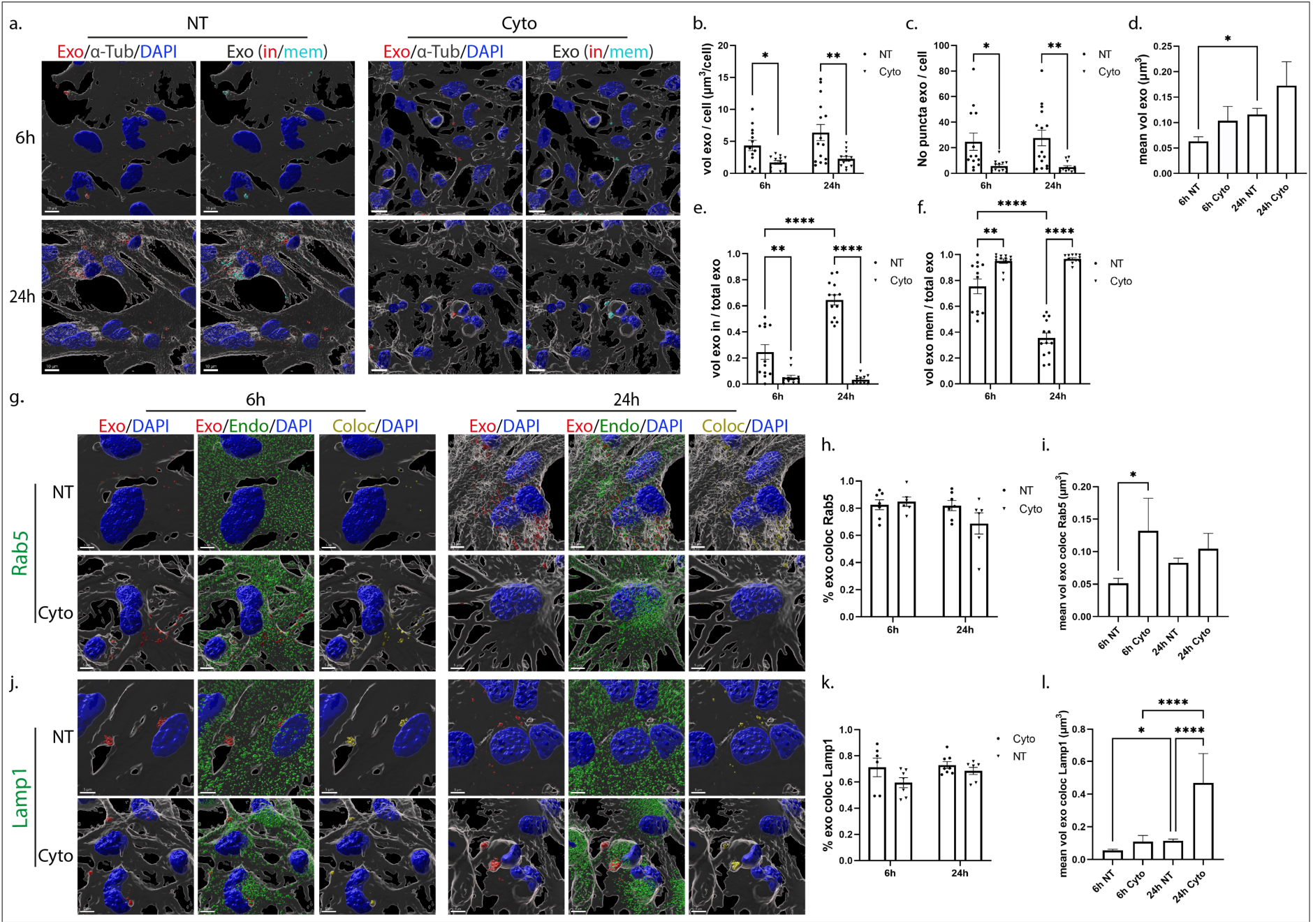
Uptake of exosomes in primary astrocytes through the actin-dependent endocytic pathway. Cells, incubated with Dil-labeled exosomes (red), in the absence or presence of Cytochalasin D (Cyto) for 6h and 24h, were fixed and immunolabeled against α-Tubulin (α-Tub) (gray) and nuclei were stained with DAPI (blue). Representative Imaris images show the internalization of exosomes and in/mem exosomes without or with Cyto, 6h and 24h (a) post-treatment. Scale bar 10 μm. Graphs present the total volume of internalized exosomes per cell (b), the number of exosomal puncta per cell (c), the mean volume of internalized exosomes (d), the percentage of exosomal volume, inside (e) or membrane (f) per total volume, 6h and 24h post-incubation. Colocalization of exosomes with Rab5 (g-i) and Lamp1 (j-l) was evaluated. Scale bar 5 μm. Graphs show colocalization between exosomes and Rab5 (h) or Lamp1 (k) and the mean volume of colocalized puncta (i and l, respectively) at different time points, with or without Cyto. Data are presented as the mean ± SEM of minimum 3 independent cell preparations, with at least two replicates per assay; one-way ANOVA with Tukey’s correction was used for (d), (i) and (l), two-way ANOVA with Tukey’s correction for (e), (f), (h) and (k) and multiple t-test for (b) and (c). Statistical significance was set as *p < .05, **p < .01, ***p < .001, ****p < .0001.

### 3.6. Exosome-mediated transmission of fibrillar α-Synuclein in primary microglia cells

To elucidate the role of exosomes in the transmission and clearance of pathogenic protein aggregates, we used structurally well-defined human recombinant α-Syn fibrils (PFFs) that induce seeding after take up by neuronal cells (Bousset et al., 2013; Pantazopoulou et al., 2021) and monitored how exosome-association affected their internalization and clearance pathways in microglia. Recent studies have reported that exosomes associate with fibrillar α-Syn and A*β* and either block α-Syn internalization in primary neuronal cells (Karampetsou et al., 2020) or enhance A*β* clearance by microglia (Yuyama et al., 2012). We preincubated exosomes and α-Syn PFFs for 18h and exposed microglia to the exosome-associated PFFs for 2h. α-Syn internalization and intracellular trafficking, was assessed 2h and 6h post-addition. Cultures treated with α-Syn PFFs or exosomes alone were used as controls. Staining against α-Syn with the specific antibody D10 that recognises human α-Syn C-terminal end, followed by confocal microscopy and Imaris analysis, revealed that α-Syn PFFs entered microglia cells either alone or in association with exosomes (PFF+Exo). PFFs alone were taken up more efficiently than PFF+Exo by microglia, 2h post-incubation (Figure 10a,b). Interestingly, α-Syn-PFFs were cleared from the cells, whether associated or not to exosomes, 6h post-addition (Figure 10a,b). However, we observed an increase in the degradation rate of PFFs when cells were incubated with PFF-associated exosomes; 2.614 for PFFs and 3.306 for PFFs+Exo. Additionally, the size of α-Syn puncta observed inside cells was considerably higher when cells were incubated with exosome associated-PFFs (Figure 10c). Exosome association with PFFs did not affect the internalization of exosomes by microglia nor did it affect their size (Figure 10a,d,e). Altogether, our data indicate either that α-Syn PFFs are internalized faster in microglia when they are not associated with exosomes, or that α-Syn PFFs in association with exosomes are targeted more efficiently to the degradation pathways, as implied from the different clearance rates.

**Figure 10.**
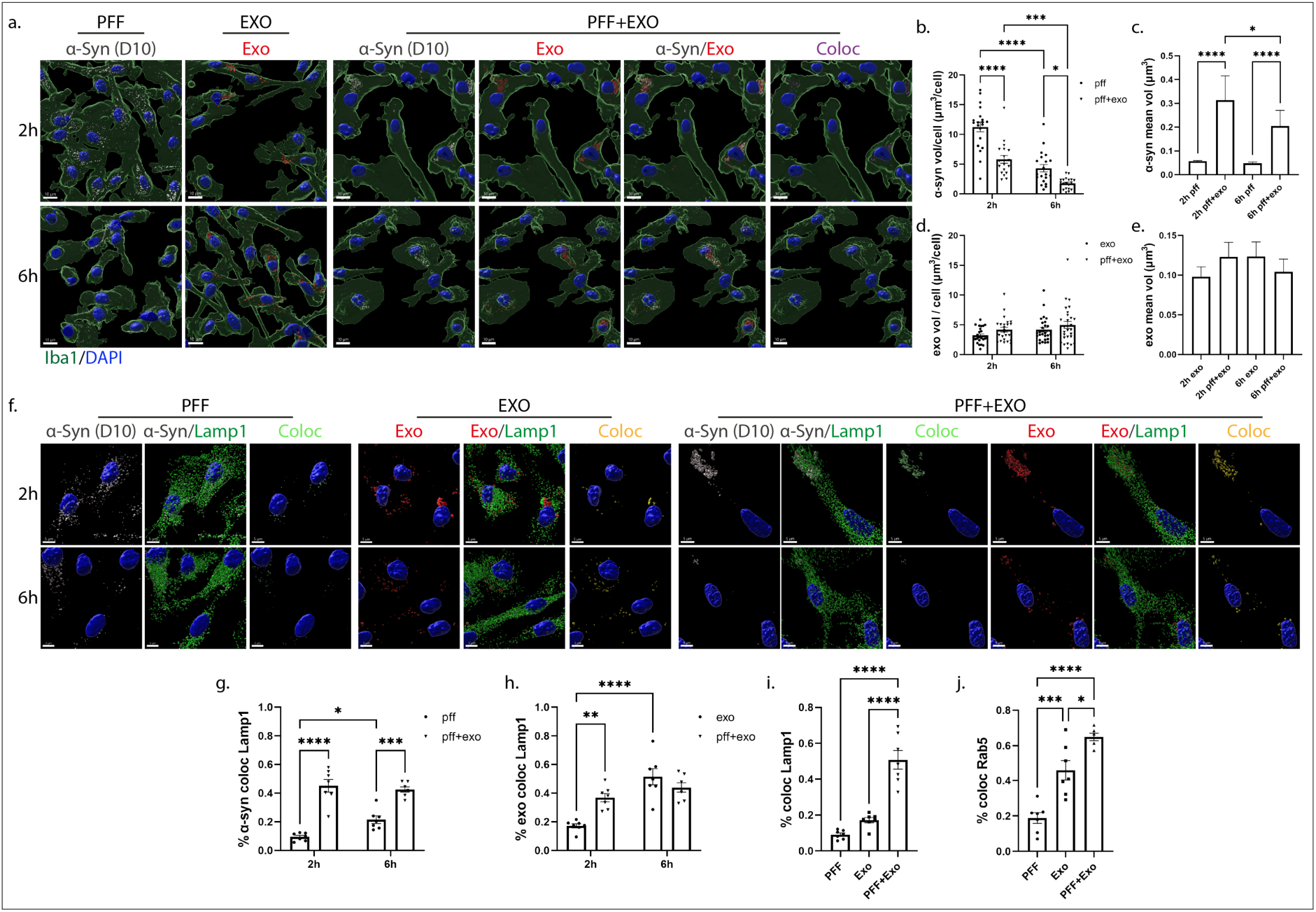
Exosome-dependent α-Syn transmission in primary microglia. α-Syn pre-formed fibrils (PFF) were pre-incubated with exosomes derived from SNCA KO mouse brains (PFF+Exo), for 18h at 37 °C. Microglia cells were treated with PFFs or EXO alone or PFF+Exo for 2h. Uptake and intracellular trafficking of α-Syn and exosomes were monitored 2h and 6h post-addition. Cells were fixed and immunostained against α-Syn (D10, light grey) and Iba1 (green). Cell nuclei were stained with DAPI (blue). Representative Imaris images show the uptake of α-Syn and exosomes in all three conditions (PFFs, EXO and PFF+Exo) 2h and 6h post-treatment. Colocalization of α-Syn with exosomes is depicted in magenta. Scale bar 10 μm. Graphs present the total volume of α-Syn per cell (b), the mean volume of α-Syn individual puncta (c), the total volume of exosomes per cell (d) and the mean volume of exosomal puncta (e) under the different conditions. The endocytic trafficking was monitored by measuring colocalization of α-Syn (depicted in light green) and exosomes (depicted in yellow) with Lamp1 (f-j), at the different treatments (PFFs, EXO and PFF+Exo). Scale bar 5 μm. Graphs show colocalization of α-Syn with Lamp1 (g) and of exosomes with Lamp1 (h) in the different conditions (PFFs, EXO, PFF+Exo), as well as colocalization of α-Syn/exo colocalized puncta with Lamp1 and Rab5, respectively, (i) and (j), in the PFF+EXO compared to the PFF or EXO alone treatments, 2h post-incubation. Data are presented as the mean ± SEM of minimum 3 independent cell preparations, with at least two replicates per assay; one-way ANOVA with Tukey’s correction was used for (c), (e), (i) and (j), two-way ANOVA with Tukey’s correction for (d), (g) and (h) and multiple t-test for (b). Statistical significance was set as *p < .05, **p < .01, ***p < .001, ****p < .0001.

To further examine this hypothesis, we examined the intracellular pathway that α-Syn PFFs followed in the absence or presence of exosomes. Staining with Rab5 and Lamp1, and following colocalization analysis with the Imaris Imaging software, we found that PFFs colocalized with both endocytic markers. However, colocalization was substantially higher in the presence of exosomes (Figure 10, Sup. Fig. 8). More specifically, in PFF-treated cells, the percentage of colocalization with Rab5 and Lamp1 was 20% and 8%, respectively, after 2h, increasing to 33% and 20%, at 6h post-PFF addition. In PFF+Exo treated cells, colocalization with both Rab5 and Lamp1 was higher (53% and 45%, respectively) after 2h of incubation, compared to PFF-alone treated cells. After 6h of treatment, colocalization with Rab5 dropped to 30% while with Lamp1 it remained at the same levels (Figure 10f,g, Sup. Fig. 8a,b). Colocalization of exosomes with Rab5 did not change following association with PFFs, on the contrary colocalization of exosomes with Lamp1 was higher when they were associated with PFFs (Figure 10f,h, Sup. Fig. 8a,c). Furthermore, α-Syn/Exo colocalized puncta displayed even higher colocalization with both Rab5 and Lamp1 (64% and 50%, respectively) when compared to cells treated with PFF-or Exo-alone, (Figure 10i,j). Our data indicate that in the presence of exosomes, fibrillar α-Syn is targeted faster to the endolysosomal pathway for subsequent degradation, suggesting a critical role of exosomes in the clearance process of α-Syn aggregates in microglia cells.

## Discussion

In the present study, we explored the internalization processes and endocytic trafficking of brain-derived exosomes in primary microglia and astrocytes. Our data indicate that brain-derived exosomes are taken up by both glial cell types, however microglia demonstrate a more efficient internalization rate compared to astrocytes (Figure 1, Sup. Fig. 9a). Both cell types utilize the actin-dependent pathways, macropinocytosis and/or phagocytosis, for the transfer of exosomes within the cells (Figure 8, 9) and subsequently target them to the endolysosomal pathway for further processing (Figure 2, 3, Sup. Fig. 9a). Additionally, we assessed the role of lipid rafts in exosome uptake (Figure 4-7) and interestingly found that exosomes are internalized faster when cholesterol is stripped off the plasma membrane in both microglia and astrocytes (Figure 6,7). Last, we investigated the potency of exosomes in sequestering pathogenic proteins, present in the extracellular space, into microglia using α-syn PFFs as a paradigm. We showed that fibrillar α-Syn species when pre-incubated with exosomes enter the endosomal pathway and are targeted to the lysosome for subsequent degradation. In the absence of exosomes, only a small portion of fibrillar α-Syn is sorted to the endosomal pathway, thus exhibiting a delay in the degradation process (Figure 10, Sup. Fig. 9b).

Exosomes mediate communication between neuron and glial cells, hence participating in various functions of the brain, both in health and disease (Pascual et al., 2020). Thus far, studies have focused mostly on glia-derived exosomes and their role in neuronal function, and little is known about exosome internalization by glial cells and their subsequent regulation. In our study, microglia uptake exosomes within 2h, and further after 6h, whereas in astrocytes large exosomal puncta reside on the plasma membrane 6h post-incubation reaching the cytoplasm only after 24h. This result indicates a slow internalization rate in astrocytes compared to microglia, the professional phagocytes of the CNS (Figure 1, 10). Under physiological conditions, exosome uptake regulates many functions of astrocytes, such as maintenance of BBB integrity, synaptic regulation and glutamate level modulation, providing neuroprotective services to neurons (Vasile et al., 2017; Ogaki et al., 2021). Microglia achieve excitatory neurotransmission and inflammatory response through exosome circulation (Paolicelli et al., 2019). Exosome uptake by astrocytes may have a negative role in glioma cell invasion and tumour migration (Bian et al., 2019; Zamani Esmati et al., 2021), while after ischemic stroke exosome internalization has been shown to confer neuroprotection (Pan et al., 2016; Deng et al., 2019; Sun et al., 2019). Exosome internalization by microglia leads to secretion of proinflammatory cytokines in autism (Tsilioni and Theoharides, 2018) and triggers inflammatory response in neurodegenerative diseases such as Alzheimer’s disease (AD) and Amyotrophic lateral sclerosis (ALS) (Pinto et al., 2017; Fernandes et al., 2018). Our data suggest that microglia possess mechanisms for rapid response to extracellular changes for targeted degradation and the subsequent inflammatory response, whereas astrocytes slowly endocytose exosomes protecting the cell from unrestrained signalling response, reflecting the cells’ key functions in the CNS. Noteworthy, our experimental approach entails limitations regarding the cell-origin of exosomes derived from whole brain extracts. Future investigation is required to study the correlation of exosome origin and recipient cell specificity, mainly in astrocytes where uptake appears to be regulated.

Over the last decade, numerous reports have aimed to elucidate the internalization and endocytic trafficking of exosomes, and hence the pathways essential for cell-to-cell communication. Clathrin-mediated endocytosis is the best molecularly defined pathway involved in the uptake of exosomes by numerous cells including endothelial, cardiomyocytes, placenta cells and cancer cell lines (Holder et al., 2016; Banizs et al., 2018; Eguchi et al., 2019; Costafreda et al., 2020; Wu et al., 2021). In several cases, it was reported that more than one pathway may independently and cooperatively contribute to the endocytic processes (Tian et al., 2014; Cheung et al., 2016; Kanno et al., 2020; Tabak et al., 2021; Ginini et al., 2022). Additionally, identifying the endocytic pathway involved remains a challenge due to the cross-reactive effect of pharmacological inhibitors (Dutta and Donaldson, 2012; Rennick et al., 2021). In this regard, dynasore is used to inhibit dynamin function, implicated in vesicle fission from the plasma membrane after internalization through the clathrin- or caveolin-mediated endocytosis, but also reduces cholesterol on the plasma membrane, thus altering lipid composition in lipid rafts in a dynamin-independent manner (Preta et al., 2015). In our experiments, dynasore treatment conferred no effect on exosome uptake by primary microglia cells (Figure 4, Sup. Fig. 5), however it increased their internalization in astrocytes compared to control conditions after 6h of incubation, as most exosomes were found to localise within the cytoplasm with a small proportion remaining on the plasma membrane (Figure 5, Sup. Fig. 5). Methyl-*β*-cyclodextrin is used to inhibit lipid raft-mediated, caveolae-dependent or independent, endocytosis by stripping off cholesterol from the plasma membrane. Several studies have demonstrated the essential role of this pathway in exosome uptake by cancer cells, T-cells, and HUVECs (Escrevente et al., 2011; Svensson et al., 2013; Cheung et al., 2016; Cerezo-Magaña et al., 2021). On the contrary, in PC12 cells, exosome uptake was increased upon methyl-*β*-cyclodextrin inhibition (Tian et al., 2014). Accordingly, in our experiments, methyl-*β*-cyclodextrin treatment appeared to induce internalization in both cell types (Figure 6, 7, Sup. Fig. 6), whereas dynasore increased uptake only in astrocytes, probably because of its effect on cholesterol distribution. Taken together, exosome uptake is negatively regulated by cholesterol in both microglia and astrocytes, possibly due to alterations in plasma membrane fluidity, accelerating uptake by other pathways. Furthermore, it has been reported that methyl-*β*-cyclodextrin may facilitate the secretion of the vesicular content (Vacca et al., 2019), hence justifying the decrease of exosomes observed 24h post-incubation in astrocytes. Considering the fast internalization processes observed in microglia, no effect was detected. Of note, cholesterol levels are decreased in AD patients, mostly in brain areas more susceptible to AD (cortex and hippocampus) (Leduc et al., 2010). Additionally, TREM2, the transmembrane receptor in microglia implicated in late-onset AD, regulates cholesterol and lipid metabolism (Nugent et al., 2020). In this sense, our data could yield interesting information concerning exosome-mediated A*β*/Tau transmission, microglia activation and disease progression in AD.

Macropinocytosis and/or phagocytosis are implicated in exosome uptake by different cell types, including professional and non-professional phagocytes (Feng et al., 2010; Fitzner et al., 2011; Nakase et al., 2015; Kamerkar et al., 2017; Samuel et al., 2018; Ogese et al., 2019). Cytochalasin D is a widely used pharmacological inhibitor that blocks both pathways, through interference with actin dynamics. Other pharmacological reagents (wortmannin and LY294002) target components shared by both pathways and while amiloride seemingly inhibits macropinocytosis with high specificity, it may affect actin as well (Lagana et al., 2000; Dutta and Donaldson, 2012). In the present study, Cytochalasin D treatment perturbed internalization of exosomes in both microglia and astrocytes, leading to the conclusion that phagocytosis and/or macropinocytosis are the main endocytic pathway(s) for exosome uptake in glial cells (Figure 8,9, Sup. Fig. 7). Our results are in agreement with past studies where oligodendroglia cell-derived exosomes are restricted from entering microglia upon amiloride treatment (Fitzner et al., 2011).

Thus far, limited studies have addressed the post-internalization fate of exosomes and subsequent endosomal trafficking (Feng et al., 2010; Nanbo et al., 2013; Li et al., 2020). Exosomes, internalized by glial cells, enter the endolysosomal pathway, localizing to Rab5- and Lamp1-positive compartments (Figure 2,3, Sup. Fig. 3,4). The differences observed in the proportion of exosomes residing in EE and LE/Lysosomes as well as in the size of colocalized exosomes between the cell types at different time points, support faster internalization processes and subcellular trafficking to lysosomes in microglia. Astrocytes exhibit a pronounced localization of exosomes at EE at early time points, as internalization is gradual, with most exosomes residing on the plasma membrane. In agreement with recent reports which detect Lamp1 in EE-positive compartments (Cook et al., 2004; Cheng et al., 2018), our data also show that exosomes residing on the plasma membrane, at early time points (6h) in astrocytic cells (Figure 3) and following Cytochalasin D treatment in microglia and astrocytes, are Lamp1-positive (Figure 8, 9). The role of Lamp1 in the plasma membrane remains elusive and further investigation linking its function with EE, LE or lysosomes is required. Interestingly, and in accordance with previous studies (Preta et al., 2015), dynasore seems to interfere with endosomal maturation and trafficking, resulting in exosomal puncta restricted from sorting to the endolysosomal pathway (Figure 4,5, Sup. Fig. 5). Cholesterol depletion is proposed to affect endosomal flux and LE maturation (Boura et al., 2012; Glitscher and Hildt, 2021), though it appeared to have a differential impact on exosome internalization by the two cell types. In methyl-*β*-cyclodextrin treated microglia, exosomes seem to be blocked in an intermediate compartment positive for both Rab5 and Lamp1 (Figure 6), whereas in astrocytic cells, exosomes probably bypass the endosomal pathway and localize in Lamp1-positive compartments suggesting their direct fusion with lysosomes (Figure 7). Future investigation may address the interplay between endosomal recycling to the plasma membrane and lysosomal sorting, elucidating cholesterol’s contribution to the distinct pathways and thus to the fate of exosomal cargo.

To further investigate the role of exosomes in the clearance of pathogenic protein aggregates by microglia, we assessed fibrillar α-Syn clearance in the presence or absence of exosomes. It has been shown that microglia cell exposure to PFFs induces the release of α-Syn-containing exosomes and pro-inflammatory cytokines, further contributing to the progression of α-Syn pathology in neurons (Guo et al., 2020). Our data demonstrate that fibrillar α-Syn fails to enter the endosomal pathway, accumulates in the cytoplasm and is cleared from the cells at later time points, possibly through autophagy, as previously described (Choi et al., 2020). Exosomes significantly facilitate the process of sorting of α-Syn PFFs to the endosomal pathway for rapid degradation by the lysosome (Figure 10, Sup. Fig. 8, 9b), as previously shown in an AD model where A*β* associated exosomes were sorted to lysosomes for subsequent degradation in microglia (Yuyama et al., 2012). Collectively, it appears that association of fibrillar α-Syn with exosomes regulates the subsequent clearance pathway followed, thus modulating degradation efficiency and likely the following inflammatory response. Although it has been suggested that the lysosome is the final destination for exosomes (Heusermann et al., 2016; Roberts-Dalton et al., 2017; Hunter et al., 2019), recent reports support the idea that their cargo can escape endosomes and be targeted to the cytoplasm where they can exert their function (Flavin et al., 2017; Li et al., 2020; Bonsergent et al., 2021; Polanco et al., 2021). This is a proposed mechanism through which exosome-delivered pathogenic proteins escape lysosomal degradation through vesicular rupture and initiate pathogenic processes in the cytoplasm. Thus, to further examine the role of exosomes in α-Syn transmission and propagation, it would be interesting to investigate fibrillar α-Syn-induced vesicular rupture and/or exosome release in our model, as well as the effect on microglia activation and inflammatory response, and in turn transmission and disease progression in neurons.

For our study, we utilized high-resolution confocal images and through 3D reconstruction with the Imaris Imaging software, we thoroughly analysed exosome internalization and endosomal sorting using a variety of parameters, rigorously dissecting the pathways involved. To the best of our knowledge, this is the first report that characterizes the endocytic processes implicated in exosome uptake by astrocytes. Noteworthy, we provide information on the endosomal sorting pathways implicated in exosome subcellular trafficking in both glial cell types, a field yet understudied in the CNS, and ultimately designate the differences between the cell types, hence the distinct responses of glial cells in processing exosomes and their cargo, possibly reflecting their function in the synapse. Furthermore, our results could prove to be a valuable tool in future research addressing propagation and clearance of exosome-transmitted pathogenic protein aggregates in neurodegenerative diseases, so that rapid degradation could be achieved, and inflammatory responses be modified through manipulation of endocytic pathways and intracellular trafficking to endo-lysosomes. Collectively, our study delineates important pathways involved in exosome trafficking in the CNS, paving the way for a better understanding of cell-to-cell communication in health and disease.

## Abbreviations

AD: Alzheimer’s disease
ALS: amyotrophic lateral sclerosis
α-Tub: α-Tubulin
α-Syn: α-Synuclein
BBB: blood brain barrier
CDE: caveolin-dependent endocytosis
CME: clathrin-mediated endocytosis
CNS: central nervous system
Cyclo: methyl-β-cyclodextrin
Cyto: cytochalasin D
Dyn: dynasore
EE: early endosomes
EM: electron microscopy
EV: extracellular vesicles
Exo: exosome
Exo in: exosome inside
Exo mem: exosome membrane
LE: late endosomes
PD: Parkinson’s disease
PFF: pre-formed fibrils.

## Acknowledgements

We thank Maria-Eirini Bafaloukou for preliminary results in exosome uptake by microglia. This work was co-financed by Greece and the European Union (European Social Fund-ESF) through the Operational Programme “Human Resources Development, Education and Lifelong Learning 2014–2020” (MIS 5049249). This project also received funding from the Innovative Medicines Initiative 2 Joint Undertaking under grant agreement No 116060 (IMPRiND) to KV, LS and RM. This Joint Undertaking receives support from the European Union’s Horizon 2020 research and innovation programme and EFPIA. This work is supported by the Swiss State Secretariat for Education, Research and Innovation (SERI) under contract number 17.00038. RM was further supported by France Parkinson. The opinions expressed and arguments employed herein do not necessarily reflect the official views of these funding bodies.

## Competing interests

The authors declare that they have no competing interests.

## Author contributions

MP conceived, performed experiments and analysed data, prepared figures and co-wrote the manuscript; AA performed and analysed experiments, and co-wrote the manuscript; AL conceived and performed experiments; AD and SNP contributed to imaging and statistical analysis; RM supervised and AC produced and characterized α-Syn assemblies; LS analysed experiments and co-wrote the manuscript; KV conceived and analysed experiments, and co-wrote the manuscript. All authors contributed to proof editing and finalizing the manuscript.

## Notes

### Competing Interest Statement

The authors have declared no competing interest.

