## Supporting Information for "Microglia and astrocytes differentially endocytose exosomes facilitating alpha-Synuclein endolysosomal sorting"

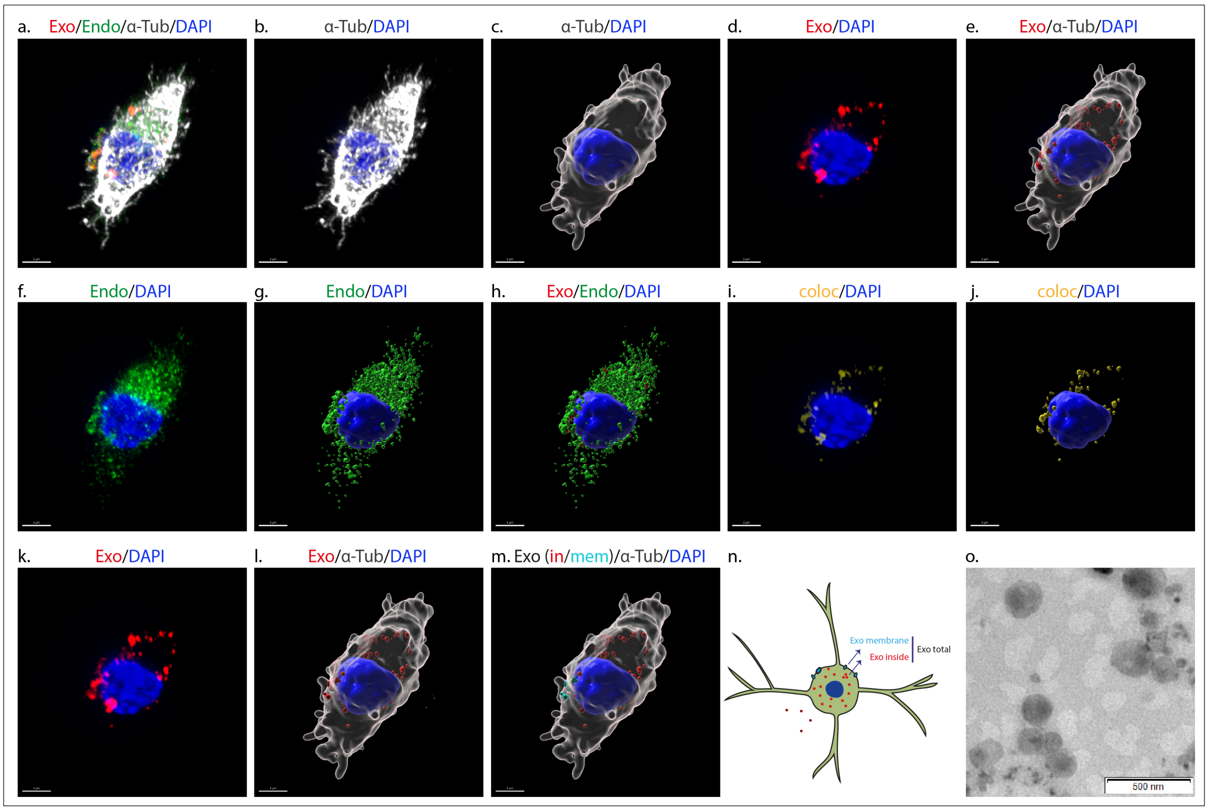


**Sup. Fig. 1. Analysis of fluorescent confocal images with the Imaris Imaging Software.** Confocal channels after deconvolution were imported into the Imaris Software (a). From the α-Tubulin (gray) and DAPI (blue) channels (b), corresponding Imaris surfaces were created (c). To measure total exosomes per cell, puncta of exosomes per cell and the mean volume of the individual puncta, α-Tubulin masked channels and surfaces corresponding to exosomes were created (d and e). To measure colocalization of exosomes with Rab5 and Lamp1, first the same procedure was followed for Rab5 and Lamp1; corresponding α-Tubulin-masked channels were created, (f), and converted into surfaces (g). The fluorescent signal from the exosome, Rab5 and Lamp1 surfaces, (h), was isolated into new masked channels that were imported into the "Coloc" module of Imaris in order to build the colocalization channel (i) and respective surface (yellow) (j) for each pair of markers and evaluate the Manders' coefficients of interest. In order to analyze the volume of internalized exosomes (the α-Tubulin-masked exosome surface already created consists of fully internalized exosomes and the fractions of membranous exosomal puncta lying within the α-Tubulin surface; here we are only interested in the former), a surface based on the original exosome channel (red, k), was created, (l), and Imaris' ''Distance transformation'' function was used to compute the distance of its objects (exosomal puncta) from the ''α-Tubulin'' surface. Zero-valued puncta (''exo total'') in the resulting "Minimum distance" statistic correspond to exosomes that have entered the cell entirely ("exo inside") and those located on the plasma membrane and are in the process of entering ("exo membrane"), (m, red and cyan blue, respectively, and in schematic illustration, n). The "exo inside" puncta are characterized by being zero-valued in the "Maximum distance" statistic, also created from the "Distance transformation" function. Scale bar 3 μm. Schematic illustration of exo total, exo in and exo mem (n). Negatively stained TEM of exosomes. Scale bar 500 nm (o).

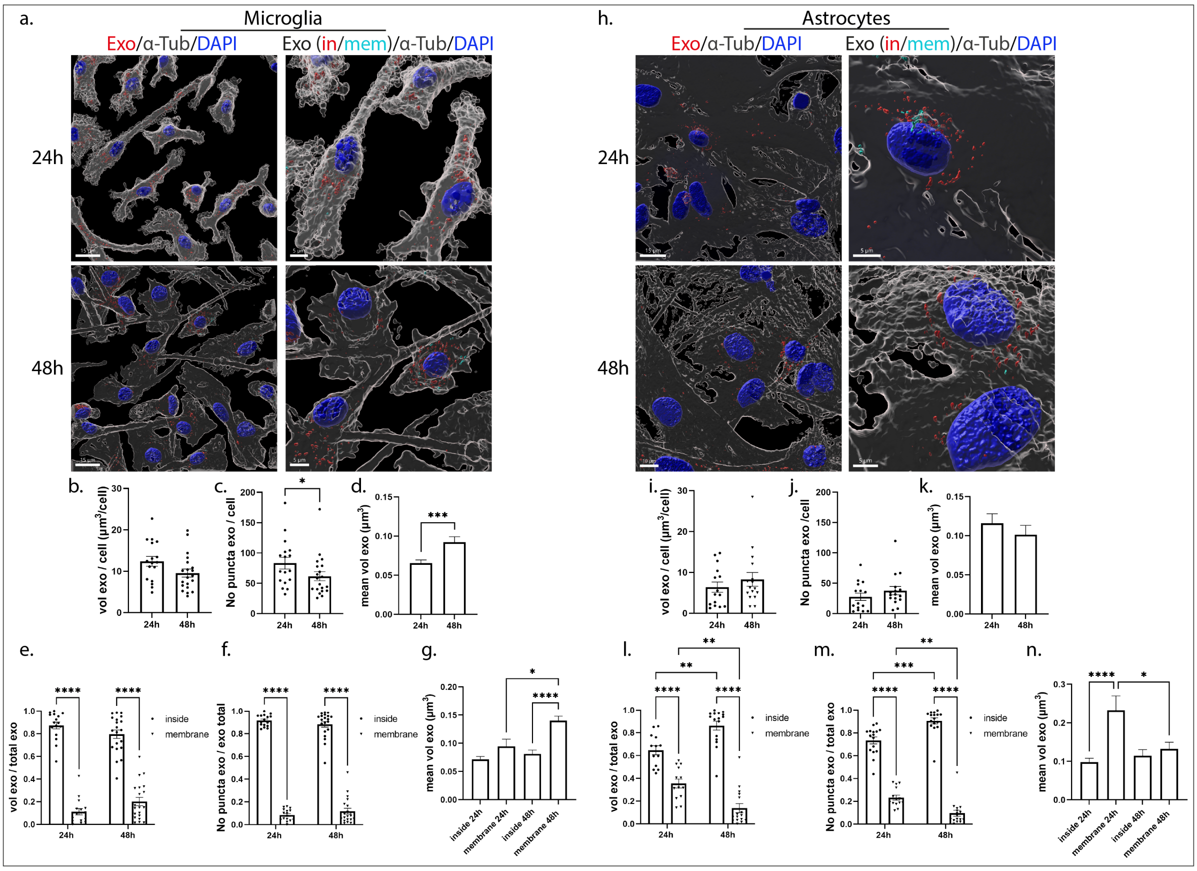


**Sup. Fig. 2. Endocytosis of brain-derived exosomes in primary microglia and astrocytes 24h and 48h post-treatment.** Primary cells were incubated with Dil-stained exosomes (depicted in red) for 24h, were washed, and internalization of exosomes was monitored at 24h and 48h of treatment. Cells were fixed and immunostained with an antibody against α-Tubulin (α-Tub) (gray) while cell nuclei were stained with DAPI (blue). Confocal images were deconvolved and analyzed with the Imaris Imaging software while exosomes were compartmentalized as cytoplasmic (exo inside, red) or membranous (exo membrane, cyan blue). Representative Imaris images depict the trafficking of exosomes masked with the tubulin surface (left panel, scale bar 15 μm) and in/mem exosomes (right panel, scale bar 5 μm) in microglia (a-g) and astrocytes (h-n) 24h and 48h post-addition. Graphs show the total volume of internalized exosomes per cell (b, i), the number of puncta per cell (c, j), the mean volume of exosomes (d, k), the ratio of the volume of exosomes (in/mem) per total volume (e, l), the ratio of the number of puncta (in/mem) per total number (f, m) and the mean volume of exosomes (in/mem) (g, n), in microglia and astrocytes respectively. Data are presented as the mean ± SEM of minimum 3 independent cell preparations, with at least two replicates per assay; Student's t-test was used for (b), (d) and (k), Mann-Whitney test for (c), (i) and (j), one-way ANOVA for (g) and (n), two-way ANOVA with Tukey's correction for (e) and (l) and multiple t-test for (f) and (m). Statistical significance was set as *$p < .05$, **$p < .01$, ***$p < .001$, ****$p < .0001$.

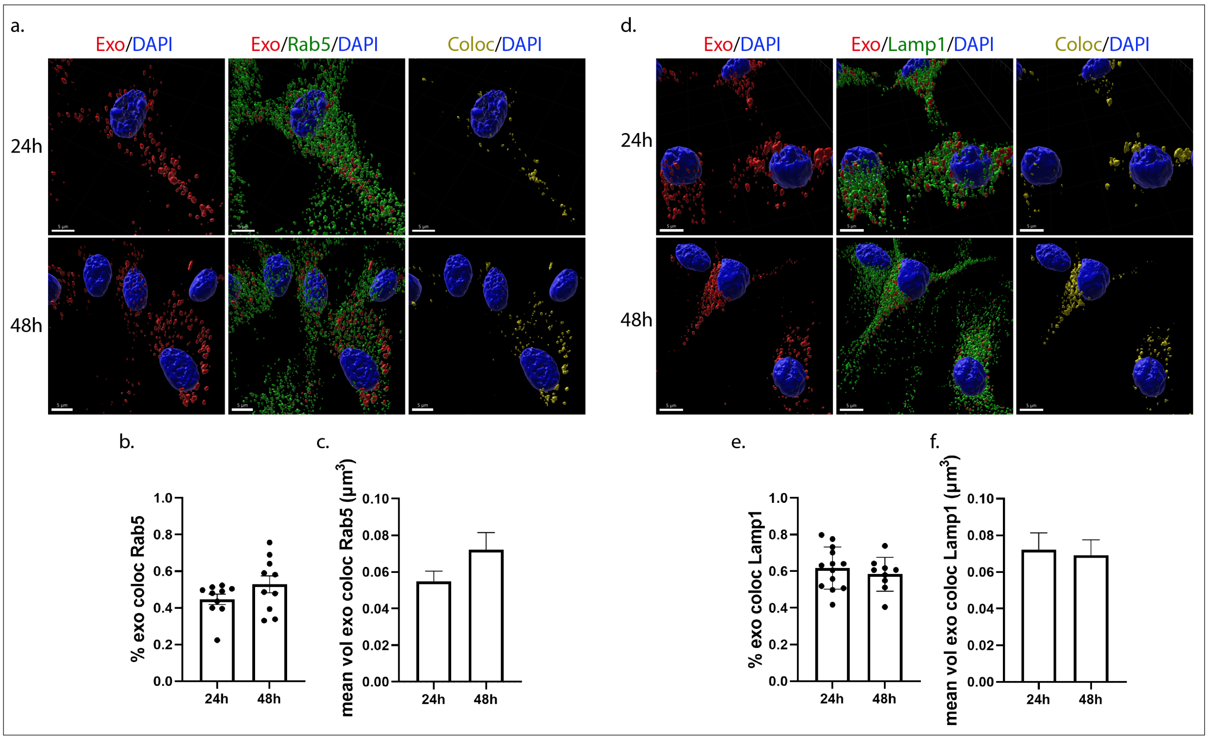


**Sup. Fig. 3. Exosomes follow the endocytic pathway and are colocalized with Rab5 (EE) and Lamp1 (LE/Lysosomes) in primary microglia, at late time points.** Cells, incubated with Dil-labeled exosomes (red) for 24h, were washed and colocalization of exosomes with Rab5 and Lamp1 was monitored 24h and 48h post-treatment. Cells were fixed and immunolabeled for Rab5 or Lamp1 (green), α-Tubulin (α-Tub) (gray) and DAPI (blue). Representative Imaris images depict colocalization between exosomes and the endocytic markers, Rab5 (a-c) and Lamp1 (d-f), 24h and 48h post-addition. Scale bar 5 μm. Graphs show colocalization between exosomes and Rab5/Lamp1 (Manders' Colocalization Coefficient) after 24h and 48h (b and e, respectively) of treatment as well as the mean volume of puncta colocalized with Rab5 (c) and Lamp1 (f) at different time points. Data are presented as the mean ± SEM of minimum 3 independent cell preparations, with more than 80 cells measured; Student's t-test was used, and statistical significance was set as $*p < .05$, $**p < .01$, $***p < .001$, $****p < .0001$.

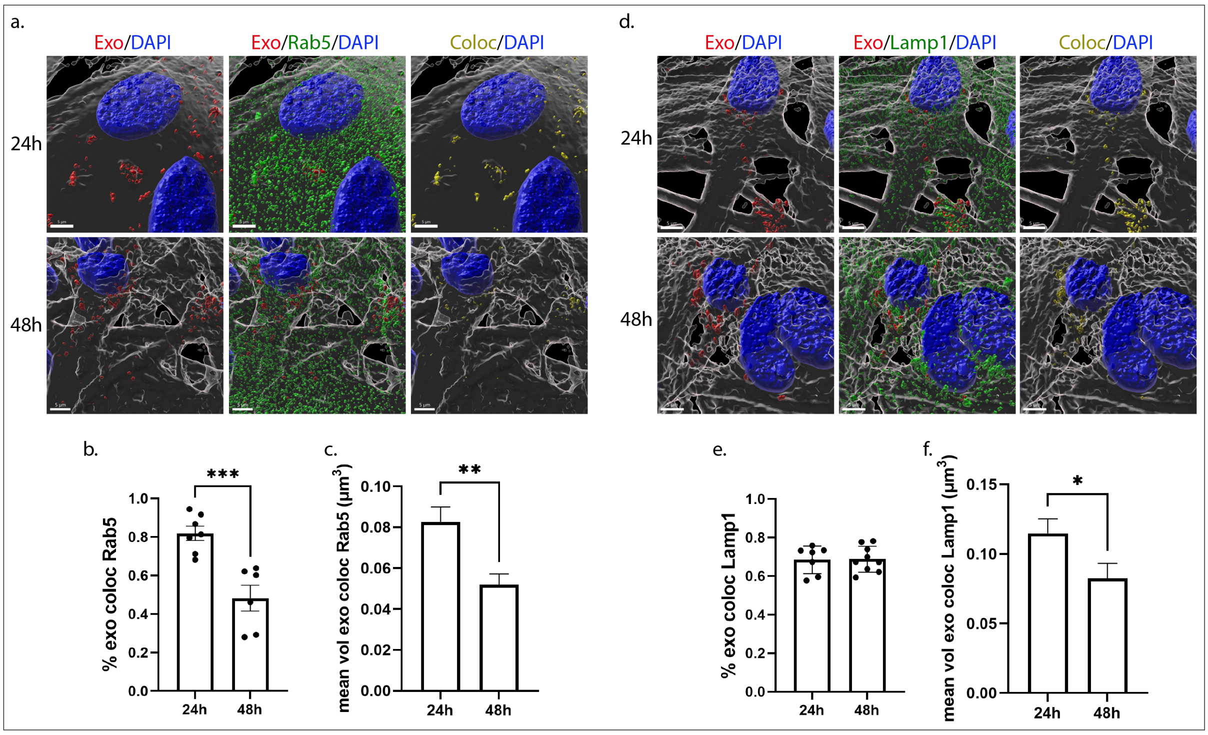


**Sup. Fig. 4. Exosomes follow the endocytic pathway and are colocalized with Rab5 (EE) and Lamp1 (LE/Lysosomes) in primary astrocytes, at late time points.** Cells, incubated with Dil-labeled exosomes (red) for 24h, were washed and the endocytic trafficking of exosomes was monitored 24h and 48h post-treatment. Cells were fixed and immunolabeled for Rab5 or Lamp1 (green), α-Tubulin (α-Tub) (gray) and DAPI (blue). Representative Imaris images depict colocalization between exosomes and the endocytic markers, Rab5 (a-c) and Lamp1 (d-f), 24h and 48h post-addition. Scale bar 5 μm. Graphs show colocalization between exosomes and Rab5/Lamp1 (Manders' Colocalization Coefficient) after 24h and 48h (b and e, respectively) of treatment as well as the mean volume of puncta colocalized with Rab5 (c) and Lamp1 (f) at different time points. Data are presented as the mean ± SEM of minimum 3 independent cell preparations, with more than 60 cells measured; Student's t-test was used, and statistical significance was set as *$p < .05$, **$p < .01$, ***$p < .001$, ****$p < .0001$.

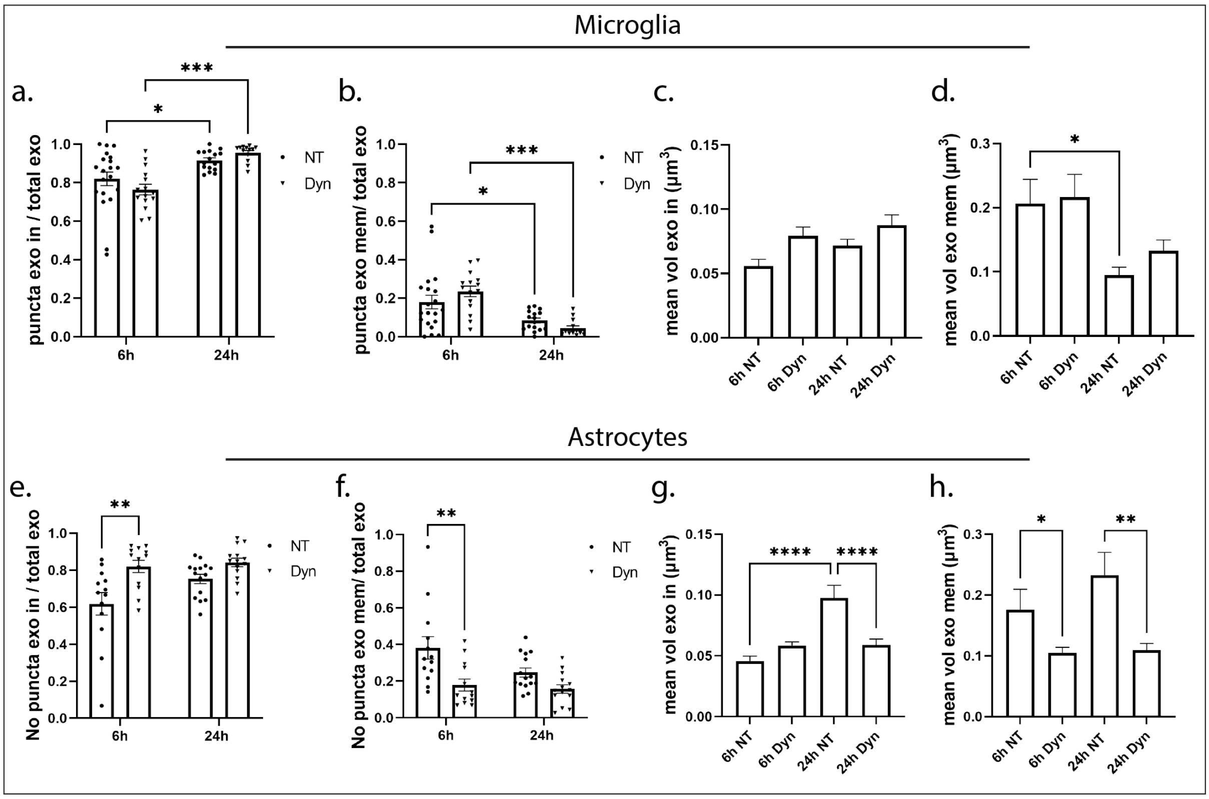


**Sup. Fig. 5. Uptake of exosomes in primary microglia and astrocytes following inhibition of the Dynamin-dependent endocytic pathway** (continued from Figure 4a and 5a)**.** Graphs show the percentage of puncta, inside (a, e) and membrane (b, f) per total puncta, and the mean volume of exosomes, inside (c, g) or membrane (d, h), after 6h and 24h of treatment, with or without dynasore, in microglia and astrocytes, respectively. Data are presented as the mean ± SEM of minimum 3 independent cell preparations; one-way Anova was used for (c), (d), (g) and (h) and multiple t-test for (a), (b), (e) and (f), and statistical significance was set as *p < .05, **p < .01, ***p < .001, ****p < .0001.

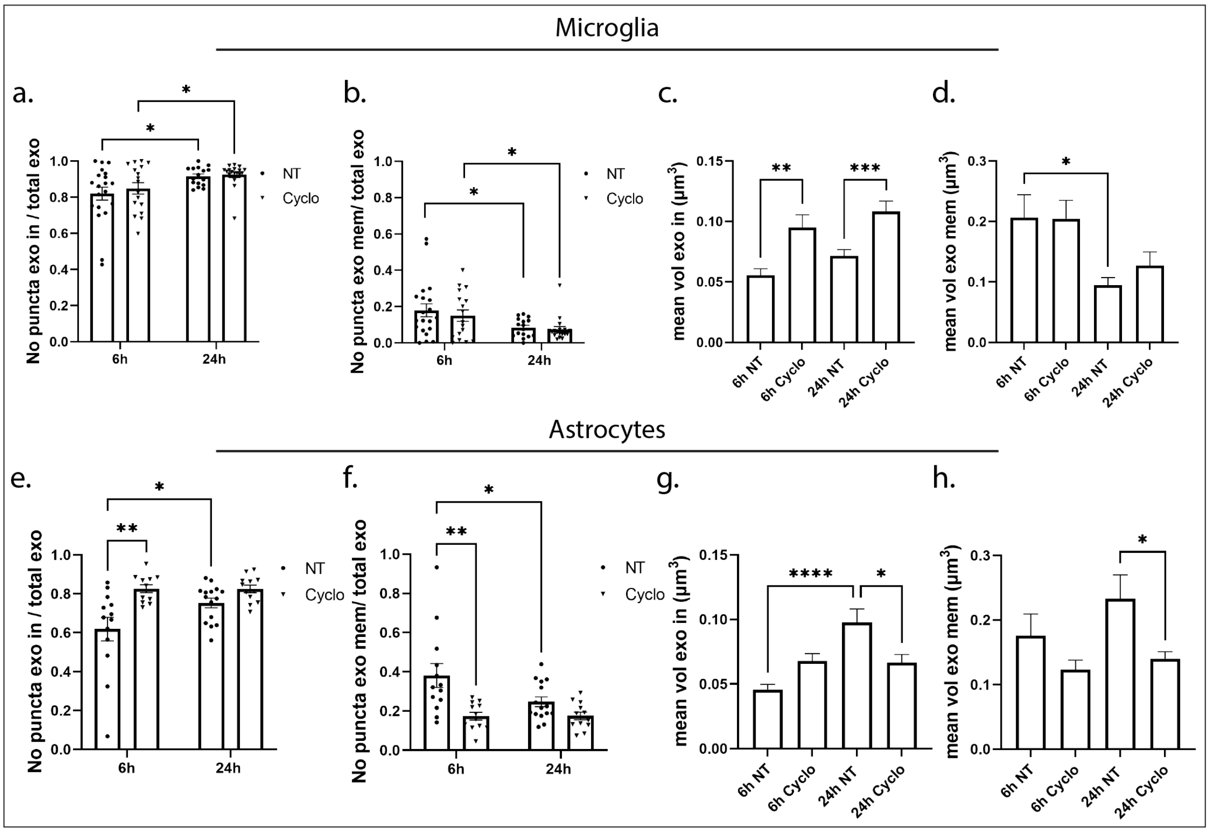


**Sup. Fig. 6. Internalization of exosomes in primary microglia and astrocytes upon inhibition of lipid raft-mediated endocytosis** (continued from Figure 6a and 7a)**.** Graphs show the percentage of puncta, inside (a and e) and membrane (b and f), per total puncta, and the mean volume of exosomes, inside (c and g) or membrane (d and h), after 6h and 24h of treatment, with or without methyl-β-cyclodextrin (cyclo), in microglia and astrocytes, respectively. Data are presented as the mean ± SEM of minimum 3 independent cell preparations; one-way Anova was used for (c), (d), (g) and (h) and multiple t-test for (a), (b), (e) and (f), and statistical significance was set as *$p < .05$, **$p < .01$, ***$p < .001$, ****$p < .0001$.

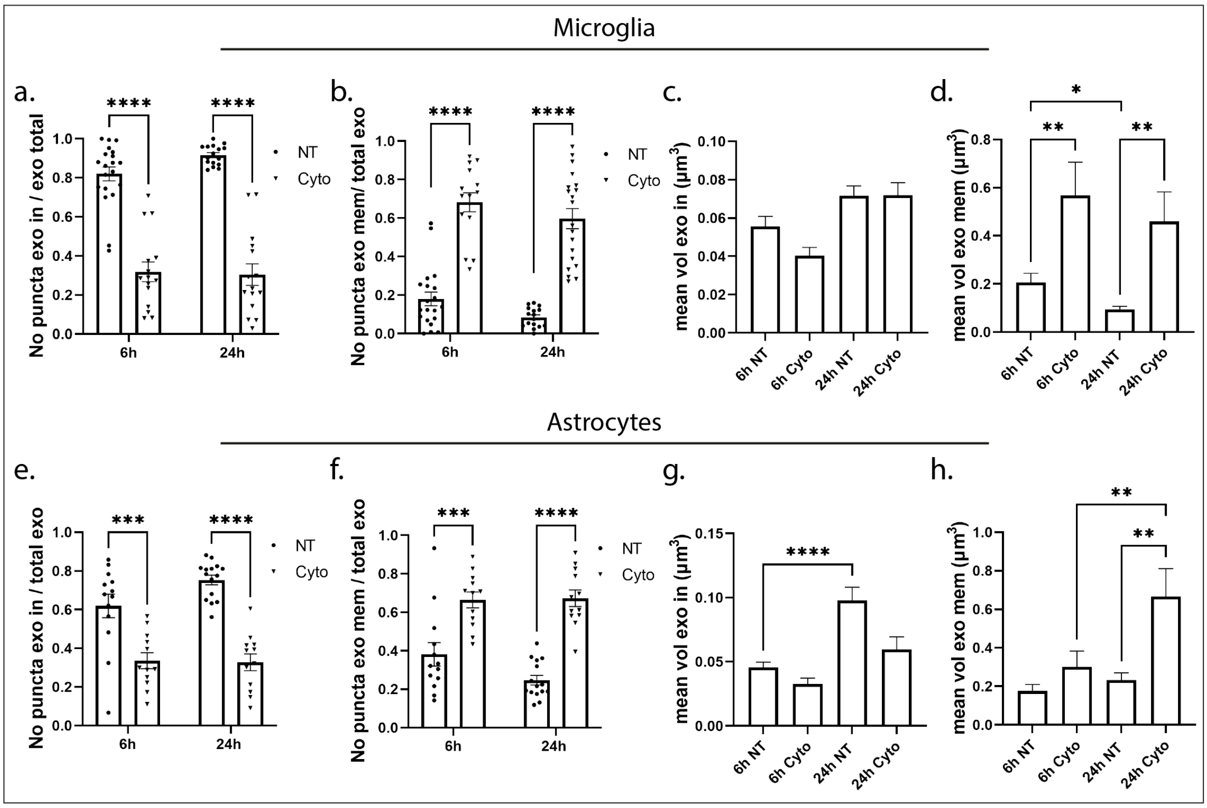


**Sup. Fig. 7. Internalization of exosomes in primary microglia and astrocytes through the actin-dependent endocytic pathway** (continued from Figure 8a and 9a**)**. Graphs show the percentage of puncta, inside (a and e) and membrane (b and f), per total puncta, and the mean volume of exosomes, inside (c and g) or membrane (d and h), after 6h and 24h of treatment, with or without Cytochalasin D (cyto), in microglia and astrocytes, respectively. Data are presented as the mean ± SEM of minimum 3 independent cell preparations; one-way Anova was used for (c), (d), (g) and (h), two- way Anova for (a) and (b), and multiple t-test for (e) and (f), and statistical significance was set as *p < .05, **p < .01, ***p < .001, ****p < .0001.

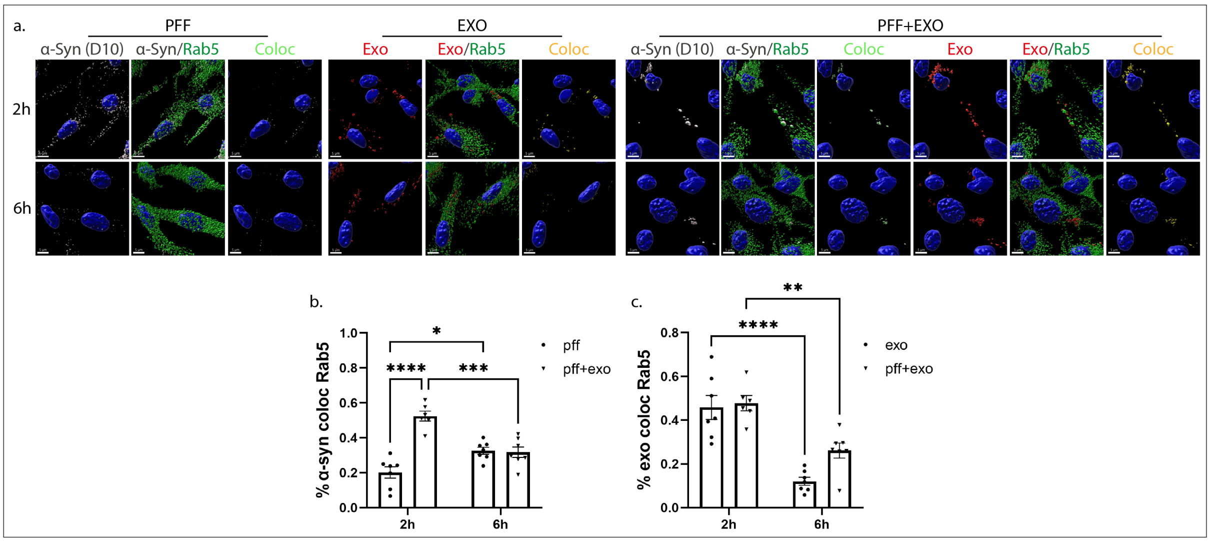


**Sup. Fig. 8. Exosome-dependent α-Syn transmission in primary microglia**. α-Syn pre-formed fibrils (PFF) were pre-incubated with exosomes derived from SNCA KO mouse brains (PFF+EXO), for 18h at 37 °C. Microglia cells were single treated with PFFs or EXO and double-treated with PFF+EXO for 2h, were washed and intracellular trafficking of α-Syn and exosomes was monitored 2h and 6h post-addition. Cells were fixed and immunostained against α-Syn (D10, light grey) and Rab5 (green). Cell nuclei were stained with DAPI (blue). The endocytic trafficking was monitored by measuring colocalization of α-Syn (depicted in light green) and exosomes (depicted in yellow) with Rab5, at the different treatments (PFFs, EXO and PFF+EXO) (a). Scale bar, 5 μm. Graphs show colocalization of α-Syn and of exosomes with Rab5 (b and c) at the different conditions (PFFs, EXO, PFF+EXO). Data are presented as the mean ± SEM of minimum 3 independent cell preparations, with at least two replicates per assay; two-way Anova was used and statistical significance was set as *$p < .05$, **$p < .01$, ***$p < .001$, ****$p < .0001$.

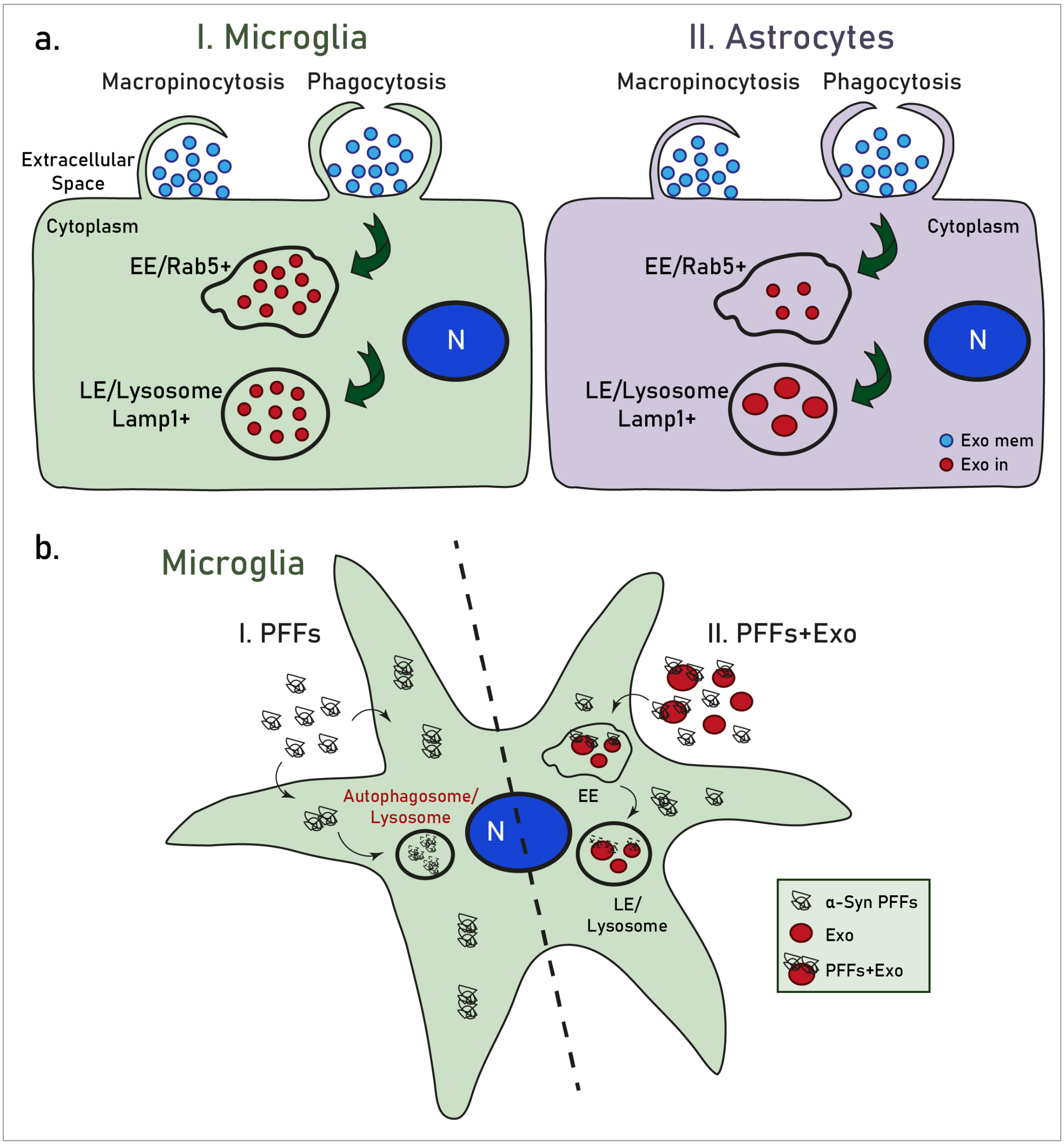


**Sup. Fig. 9. a. Internalization processes and endocytic trafficking of brain-derived exosomes in microglia (I) and astrocytes (II).** Brain-derived exosomes are taken up by both glial cell types, however microglia demonstrate a more efficient internalization rate compared to astrocytes. Both cell types utilize the actin-dependent pathways, macropinocytosis and/or phagocytosis, for the transfer of exosomes within the cells and subsequently target them to the endolysosomal pathway for further processing. **b. Exosome-dependent α-Syn transmission in microglia**. Fibrillar α-Syn-associated exosomes (PFFs+Exo) enter the endosomal pathway (EE and LE) and are targeted to the lysosome for subsequent degradation. In the absence of exosomes, α-Syn PFFs fail to enter the endosomal pathway, accumulate in the cytoplasm and are cleared from the cells at later time points, possibly through autophagy (Choi et al., 2020).
